# iPSC-Derived Myotubes from ME/CFS Patients Reveal Signature Genes, Isoforms and Pathways with Drug Repurposing Potential

**DOI:** 10.64898/2026.10.01.755863

**Authors:** Dao Nguyen, Trang T. T. Truong, Bruna Panizzutti, Megan Ellis, Briana Spolding, Courtney Swinton, Chiara Cristina Bortolasci, Jenny Bryce, Jess Smith, Cassie Field, Zoe Shih-Jung Liu, Jee Hyun Kim, Michael Berk, Ken Walder

**Affiliations:** Deakin Institute for Mental and Physical Health and Clinical Translation (Deakin IMPACT), School of Medicine, Deakin University, Australia; Florey Institute of Neuroscience and Mental Health, Parkville, Australia; Mental Health Drug and Alcohol Services, Barwon Health, University Hospital Geelong, Geelong, Australia

**Keywords:** ME/CFS, patient-derived iPSC, myotubes (MTs), DEGs, RNA sequencing, pathways, gene signature, transcriptomics

## Abstract

Myalgic encephalomyelitis/chronic fatigue syndrome (ME/CFS) is a complex multisystem disorder. Its hallmark feature, post-exertional malaise (PEM), is a delayed and prolonged worsening of symptoms following even minor exertion. As PEM commonly includes muscle pain, weakness, exercise intolerance and impaired recovery, skeletal muscle dysfunction may contribute to its pathophysiology, but the underlying molecular processes remain poorly understood. We generated induced pluripotent stem cell-derived skeletal muscle myotubes from individuals with ME/CFS (n = 6) and healthy controls (n = 7) to investigate disease-associated transcriptomic alterations and identify potential therapeutic candidates. Myogenic progenitor cells from both groups differentiated into elongated, multinucleated myotubes expressing titin and myosin heavy chain, while qPCR profiling of myogenic differentiation markers further supported successful differentiation. Gene-level analysis identified three differentially expressed genes at an FDR-adjusted *p* < 0.05: *ERVH48-1, CDX4* and *SLC2A14*. Isoform-switching analysis identified 16 genes with significant differential isoform usage, including *CACNA1H, MIEF2, SMAD2* and *CHD4*. Gene set enrichment analysis identified broader pathway-level alterations involving RNA processing, protein synthesis, proteostasis, mitochondrial quality control, metabolism and muscle-related processes. Library of Integrated Network-based Cellular Signatures Phase 2 (LINCS2) analysis further prioritised 14 compounds associated with mitochondrial metabolism, glucose and lipid regulation, PPAR signalling, calcium and excitatory signalling, neuroimmune activity and cellular stress responses. Overall, ME/CFS iPSC-derived myotubes showed modest gene-level differences alongside broader isoform- and pathway-level alterations, supporting their use for investigating skeletal muscle biology and prioritising candidates for functional validation.

## Introduction

Myalgic encephalomyelitis/chronic fatigue syndrome (ME/CFS) is a complex, disabling condition characterised by profound fatigue, post-exertional malaise (PEM), cognitive dysfunction, sleep disturbance, pain and a range of autonomic, immune and metabolic features [1–4]. PEM, defined as a delayed and prolonged worsening of symptoms following minimal physical, cognitive or emotional exertion, is a core feature of the illness and remains one of the most debilitating symptoms for patients [1, 4]. Despite its substantial personal and societal burden, the biological mechanisms underlying PEM are poorly understood, and there are currently no approved disease-modifying treatments.

Skeletal muscle symptoms, including muscle pain, weakness, exercise intolerance and prolonged recovery period after minimal exertion, are commonly reported in PEM [4–6], suggesting that impaired skeletal muscle function may contribute to disease pathophysiology. Consistent with this, studies of skeletal muscle biopsies from people with ME/CFS have reported abnormalities in mitochondrial bioenergetics, oxidative metabolism, respiratory-chain organisation and muscle-fibre morphology, with some studies also describing mitochondrial ultrastructural changes [6–9]. However, these biological findings have not yet informed a systematic pipeline for therapeutic prioritisation. Muscle biopsies are invasive and provide only a single snapshot of tissue biology, whereas blood-based studies may not adequately capture muscle-specific mechanisms. Patient-derived cellular models are therefore needed to investigate muscle-related disease biology under controlled experimental conditions and identify compounds capable of rescuing disease-relevant abnormalities.

Induced pluripotent stem cell (iPSC)-derived disease models are powerful platforms for investigating human disease mechanisms, particularly in conditions where access to affected tissues is limited and there is no validated animal model that reproduces the complex, delayed and prolonged features of PEM [10, 11]. By retaining the donor genetic background, iPSC-derived cells can capture participant-associated cellular phenotypes and provide experimentally tractable systems for studying disease pathology, modelling clinically relevant features and testing therapeutic candidates [12–17]. Such models have supported mechanistic and drug-discovery studies in other neurological disorders, including amyotrophic lateral sclerosis [13], Alzheimer’s disease [14, 15] and Parkinson’s disease [16]. When integrated with transcriptomic profiling, patient-derived cellular models can also support computational drug repurposing by comparing disease-associated gene-expression signatures with drug-induced transcriptional signatures [18, 19]. Compounds producing opposing transcriptional profiles can thereby be prioritised as candidates predicted to reverse disease-associated molecular changes[19]. Because many drug repurposing candidates have existing pharmacological, clinical or safety information, this strategy may provide a more direct route to therapeutic evaluation than conventional de novo drug discovery. Applying this approach to iPSC-derived skeletal muscle samples may therefore provide a means to identify ME/CFS-associated molecular signatures and prioritise compounds for subsequent functional testing.

In this study, we generated iPSC-derived myotubes from individuals with ME/CFS and healthy controls to investigate disease-associated skeletal muscle molecular features through gene-level differential expression, isoform-switching and pathway analyses. For computational drug repurposing, the top 100 upregulated and 100 downregulated genes from the differential-expression analysis were queried against the LINCS2 database to identify compounds predicted to reverse the ME/CFS-associated transcriptional profile. Together, this study establishes a participant-derived myotube platform for ME/CFS and identifies molecular features and candidate compounds for future functional and therapeutic investigation.

## Materials and Methods

This study received ethics approval from Barwon Health Human Research Ethics Committee (20/202) and Deakin University Human Research Ethics Committee (2021-165), Australia.

## Study design

Participants with ME/CFS and healthy controls were recruited as part of a broader iPSC disease-modelling and drug repurposing program [12]. Detailed participant recruitment, clinical assessment, demographic information, peripheral blood mononuclear cell (PBMC) isolation, iPSC generation, iPSC characterisation, myogenic progenitor (MP) differentiation and quality control are described in our previous study [20]. Briefly, PBMCs were reprogrammed into iPSCs, expanded, quality-controlled and differentiated into myogenic progenitors MPs. This study focused on terminal differentiation of these MPs into myotubes. Downstream analyses included gene-level differential expression, transcript isoform-switching analysis, pathway enrichment, and computational drug-mapping with literature-based candidate prioritisation.

## Myotube Differentiation

Myotubes from 13 participants were differentiated from iPSC-derived MP lines. For myotube differentiation, MP cells were seeded into 24-well plates pre-coated with Matrigel (Corning) and maintained in DMEM with 1000 mg/L D-Glucose (STEMCELL Technologies), supplemented with 10% MyoCult™-SF Expansion Supplement (STEMCELL Technologies). When the cultures reached 70–80% confluency, the expansion medium was replaced with MyoCult™-SF Basal Medium (STEMCELL Technologies) containing 5% MyoCult™ Differentiation Supplement (STEMCELL Technologies). Cells were maintained in differentiation medium for 10-14 days, with medium changes every 3-4 days. Myotube formation was assessed in all 13 lines on days 12–14 of differentiation by examining cell morphology using bright-field microscopy (EVOS M7000, Thermo Fisher Scientific). Myogenic differentiation was further characterised by immunocytochemistry, as previously described [20]. Expression of myosin heavy chain (MyHC), a major contractile protein of skeletal muscle, was assessed using an anti-MyHC antibody (Abcam, CAT# B37484), while titin, a structural protein of the sarcomere, was assessed using an anti-titin antibody (Invitrogen, CAT# PA5-52379). Myogenic differentiation was additionally evaluated by qPCR analysis of PAX3 (myogenic progenitor marker), MYOD1 and MYOG (myogenic regulatory factors), MYH1, ACTA1 and ACTA1 (skeletal muscle differentiation/contractile markers), as described below. Differentiated cultures were subsequently harvested for RNA extraction and sequencing.

## RNA sequencing, differential expression gene (DEG) analysis and pathway enrichment analysis

RNA sequencing, differential gene expression analysis and pathway enrichment analysis were described in detail in our previous paper [20]. Briefly, RNA was extracted from differentiated myotubes and RNA-seq libraries were prepared using the Illumina Total RNA with Ribo-Zero Plus Library Prep Kit. Sequencing was performed on the Illumina NovaSeq X platform, generating 150 bp paired -end reads (∼100 million reads per sample). Data processing, quality control, alignment to the human reference genome (GRCh38), and gene-level quantification were performed using fastp [21], HISAT2 [22], and featureCounts [23]. Normalisation and differential expression analysis were conducted using DESeq2 [24]. Surrogate variable analysis (sva package in R) [25] was applied to account for unmeasured confounders. Gene set enrichment analysis (GSEA) was performed using the clusterProfiler package in R [26] using KEGG annotations, with gene lists pre-ranked based on the sign of log fold changes multiplied by the log10-transformed p-values from the differential analysis. Pathways with an adjusted p-value <0.05 were considered enriched, and results were visualised in Cytoscape [27].

## Alternative splicing analysis

Transcriptome assembly and isoform quantification were performed using StringTie [28], enabling the reconstruction of both known and novel transcript isoforms guided by a reference annotation. FPKM (Fragments Per Kilobase of transcript per Million mapped reads) values were calculated for each transcript across all samples. Coding potential of assembled transcripts was assessed using CPAT (Coding Potential Assessment Tool) [29], applying a coding probability cutoff of 0.725 for human transcripts. Protein domains were identified using PfamScan against the Pfam-A database [30]. Additional protein characterisation tools included SignalP 6.0 for signal peptide prediction [31], DeepLoc2 for subcellular localisation prediction [32], DeepTMHMM for transmembrane domain prediction [33], and IUPred3 for intrinsically disordered region (IDR) prediction [34]. Differential transcript usage (DTU) analysis was conducted using the IsoformSwitchAnalyzeR R/Bioconductor package [35]. Genes with expression levels below 1 FPKM and those with only a single isoform were excluded from further analysis.

Isoform switching was analysed using SatuRn [36], which applies a scalable quasi-binomial generalised linear model framework for DTU analysis. Open reading frames (ORFs) were predicted, and corresponding amino acid and nucleotide sequences were extracted for downstream functional annotation. Alternative splicing events were classified as exon skipping (ES), intron retention (IR), alternative 3’ splice site (A3), alternative 5’ splice site (A5), alternative transcription start site (ATSS), alternative transcription termination site (ATTS), or mutually exclusive exons (MES). Functional consequences of isoform switching were evaluated using a differential isoform fraction (dIF) cutoff of 0.1, assessing changes in intron retention, coding potential, nonsense-mediated decay status, domain composition, and ORF sequence similarity.

## RNA Extraction and quantitative PCR

RNA was extracted using RNeasy® mini kits (Qiagen) and reverse-transcribed to produce cDNA using Maxima H Minus first strand cDNA synthesis kit (Thermo Fisher Scientific) following the manufacturer’s instructions. The Quant-iT™ OliGreen® ssDNA Assay Kit (Invitrogen) was used to quantify the cDNA concentration in each sample as per the manufacturer’s instructions. Real-time quantitative PCR (qPCR) was used to measure the expression of specific genes with primer information as provided in Supplementary file 1. The experiments were performed on a QuantStudio 3 Real-time PCR system (Thermo Fisher Scientific), using the following protocol: 95° for 7 min, 40 cycles of 95° for 30s and 60° for 1 min followed by data acquisition, 60° for 30s, 55-95°, data acquisition and 20° for 10s. Gene expression data was quantified using the ΔΔCt method normalised to the measured cDNA concentration, GAPDH expression on a pooled sample was used as qPCR plate control, plates outside 2 standard deviations from the mean were repeated.

## *In silico* identification of drug repurposing candidates

*In silico* drug repurposing was performed using the top 100 upregulated and top 100 downregulated differentially expressed genes, selected at *p* < 0.01. These gene signatures were queried against the LINCS2 database using the signatureSearch R package [18]. Compounds were prioritised according to their normalised connectivity score (NCS), with more negative scores indicating a greater predicted ability to reverse the disease-associated transcriptional profile. Only compounds with a WTCS false discovery rate below 0.05 were retained for further assessment. The highest-ranking candidates were then evaluated through a targeted literature review. Investigational agents, drugs carrying black box warnings, veterinary medicines, topical therapies, and cytotoxic chemotherapies were excluded. This stepwise filtering process was designed to identify candidates with the strongest potential for safe and clinically relevant repurposing in the target disease.

## Statistical analysis

Statistical procedures for differential gene expression, pathway enrichment and *in silico* drug-repurposing analyses are described above. qPCR results were analysed using the ΔΔCt method and normalised to cDNA concentration, then grouped according to ME/CFS or healthy control status. Data distribution was assessed using the Shapiro-Wilk test. Comparisons between ME/CFS and control groups were performed using two-tailed unpaired Welch’s *t*-tests for normally distributed data and Mann-Whitney U tests for non-normally distributed data. A *p*-value < 0.05 was considered statistically significant. Statistical analyses were performed using GraphPad Prism version 10.3.1.

## Results

### Myotube characterisations

All 13 selected cell lines successfully differentiated into myotubes, as shown by elongated multinucleated morphology and strong expression of the skeletal muscle markers titin and myosin heavy chain (MyHC). Representative images from one ME/CFS cell line and one control cell line are shown in Figure 1A, while images for all 13 differentiated myotube lines are provided in Supplementary Figure 1. qPCR analysis showed expression of myogenic differentiation markers in both ME/CFS and control myotube cultures (Figure 1B). Early myogenic markers PAX3 and MYOD1 were expressed at relatively low levels, whereas MYOG, MYH1 and particularly ACTA1 showed higher expression, consistent with progression toward a differentiated skeletal muscle phenotype. ACTA2 (SMA) was also detected, but at lower levels than ACTA1. There were no significant differences in the expression of these markers between the ME/CFS and healthy control groups, indicating comparable myogenic differentiation profiles across the two groups.

**Figure 1.**
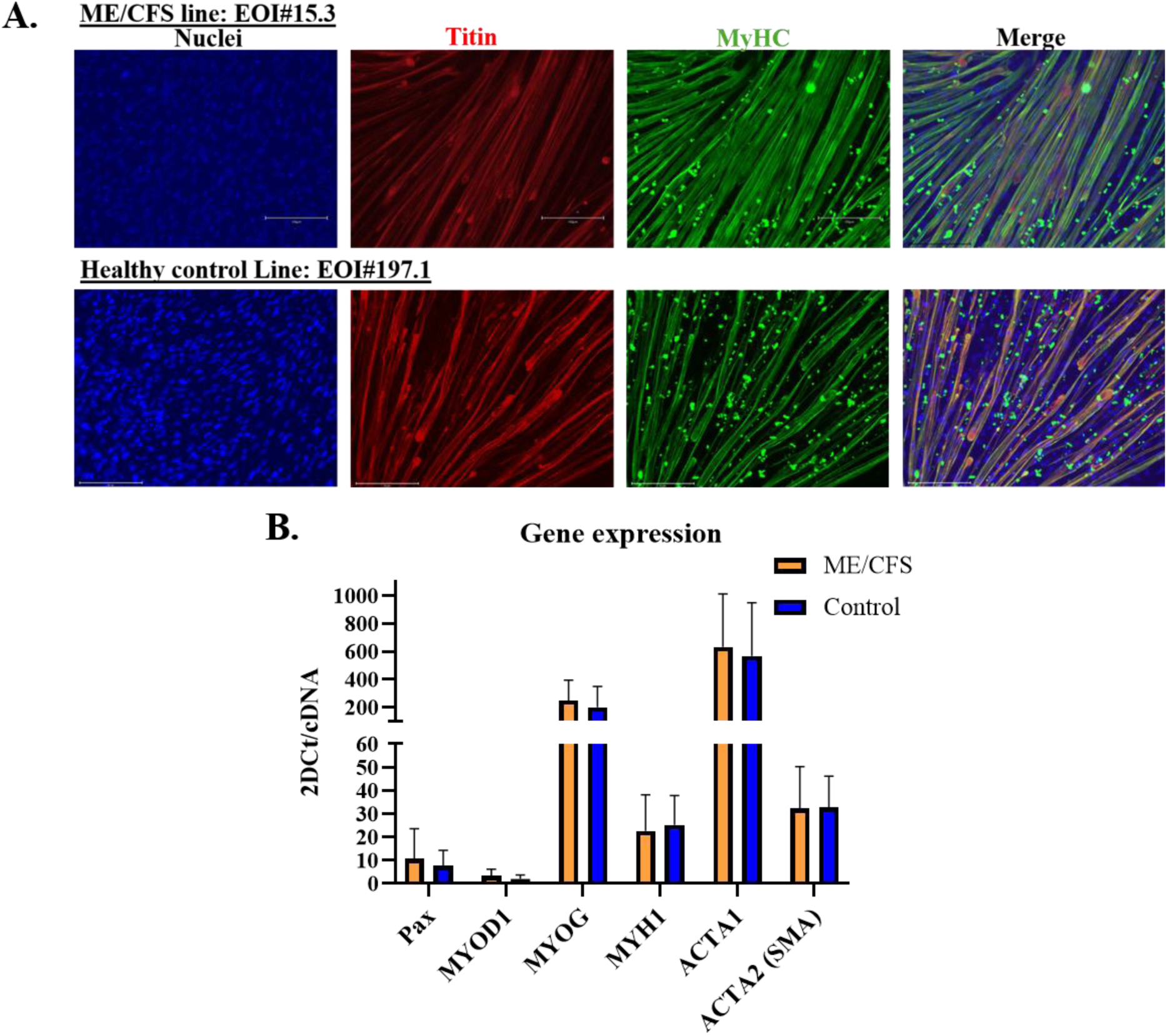
Characterisation of iPSC-derived myotubes from individuals with ME/CFS and healthy controls. Both ME/CFS and healthy control iPSC-derived myotubes exhibited elongated, multinucleated morphology and expressed the skeletal muscle markers titin and myosin heavy chain (MyHC) (**A**). qPCR analysis demonstrated expression of myogenic regulatory and skeletal muscle-associated genes (**B**). PAX3, paired box 3; MYOD1, myogenic differentiation 1; MYOG, myogenin; MYH1, myosin heavy chain 1; ACTA1, actin alpha 1, skeletal muscle; ACTA2 (α-SMA), actin alpha 2, smooth muscle. Representative images are shown; images from all differentiated lines are provided in Supplementary Figure 1.

## Differentially expressed genes and isoform usages

After normalization and false discovery rate (FDR) correction, three genes remained significantly differentially expressed between ME/CFS and control myotubes (adjusted p < 0.05): ERVH48-1, CDX4 and SLC2A14 (Table 1; Supplementary file 2). ERVH48-1, an endogenous retrovirus-related transcript, was markedly upregulated in ME/CFS myotubes and showed the largest effect size among the significant genes identified by RNA-seq (log₂ fold change = 18.27). This direction of change was confirmed by qPCR, although the magnitude of upregulation was smaller (log₂ fold change = 5.64). In contrast, two genes were significantly downregulated in ME/CFS myotubes. CDX4, a caudal-type homeobox transcription factor involved in developmental patterning and HOX-associated transcriptional programmes, was strongly downregulated by RNA-seq (log₂ fold change = -16.04), and this finding was also confirmed by qPCR (log₂ fold change = -4.17). SLC2A14, which encodes a member of the facilitated glucose transporter family, was also reduced in ME/CFS myotubes, as shown by both RNA-seq (log₂ fold change = -2.76) and qPCR (log₂ fold change = -2.05).

**Table 1.**
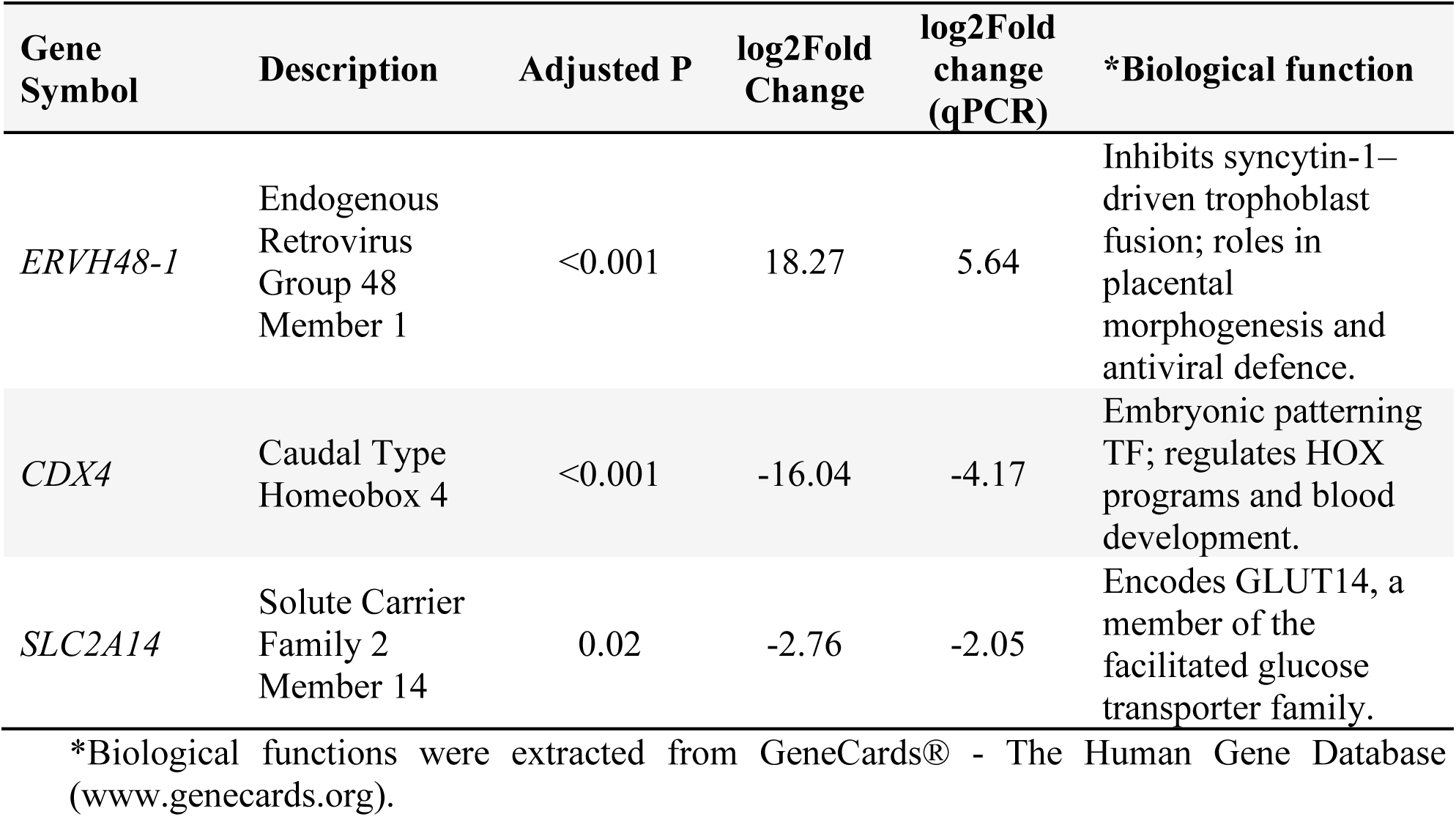
Differentially expressed genes with adjusted p < 0.05 and their functions

| Gene Symbol | Description | Adjusted P | log2Fold Change | log2Fold change (qPCR) | *Biological function |
| --- | --- | --- | --- | --- | --- |
| *ERVH48-1* | Endogenous Retrovirus Group 48 Member 1 | <0.001 | 18.27 | 5.64 | Inhibits syncytin-1–driven trophoblast fusion; roles in placental morphogenesis and antiviral defence. |
| *CDX4* | Caudal Type Homeobox 4 | <0.001 | -16.04 | -4.17 | Embryonic patterning TF; regulates HOX programs and blood development. |
| *SLC2A14* | Solute Carrier Family 2 Member 14 | 0.02 | -2.76 | -2.05 | Encodes GLUT14, a member of the facilitated glucose transporter family. |

\*Biological functions were extracted from GeneCards® - The Human Gene Database (www.genecards.org).

Given the limited number of genes that met the threshold for gene-level differential expression, we next investigated whether ME/CFS myotubes exhibited transcript-level dysregulation. This analysis identified 16 genes with significant isoform switching between ME/CFS and control cell lines (Table 2), indicating that disease-associated transcriptional changes extended beyond alterations in total mRNA abundance.

**Table 2.**
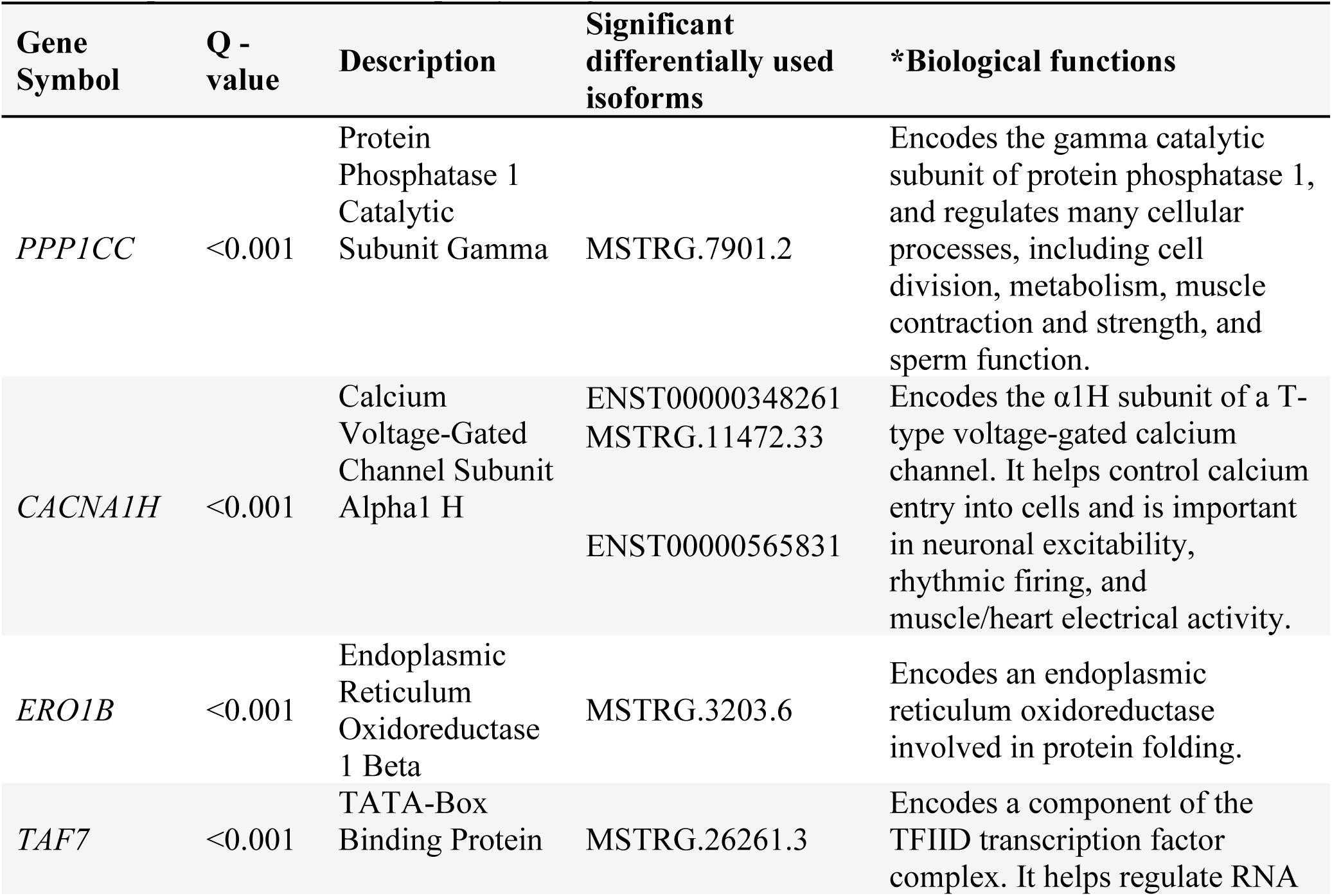
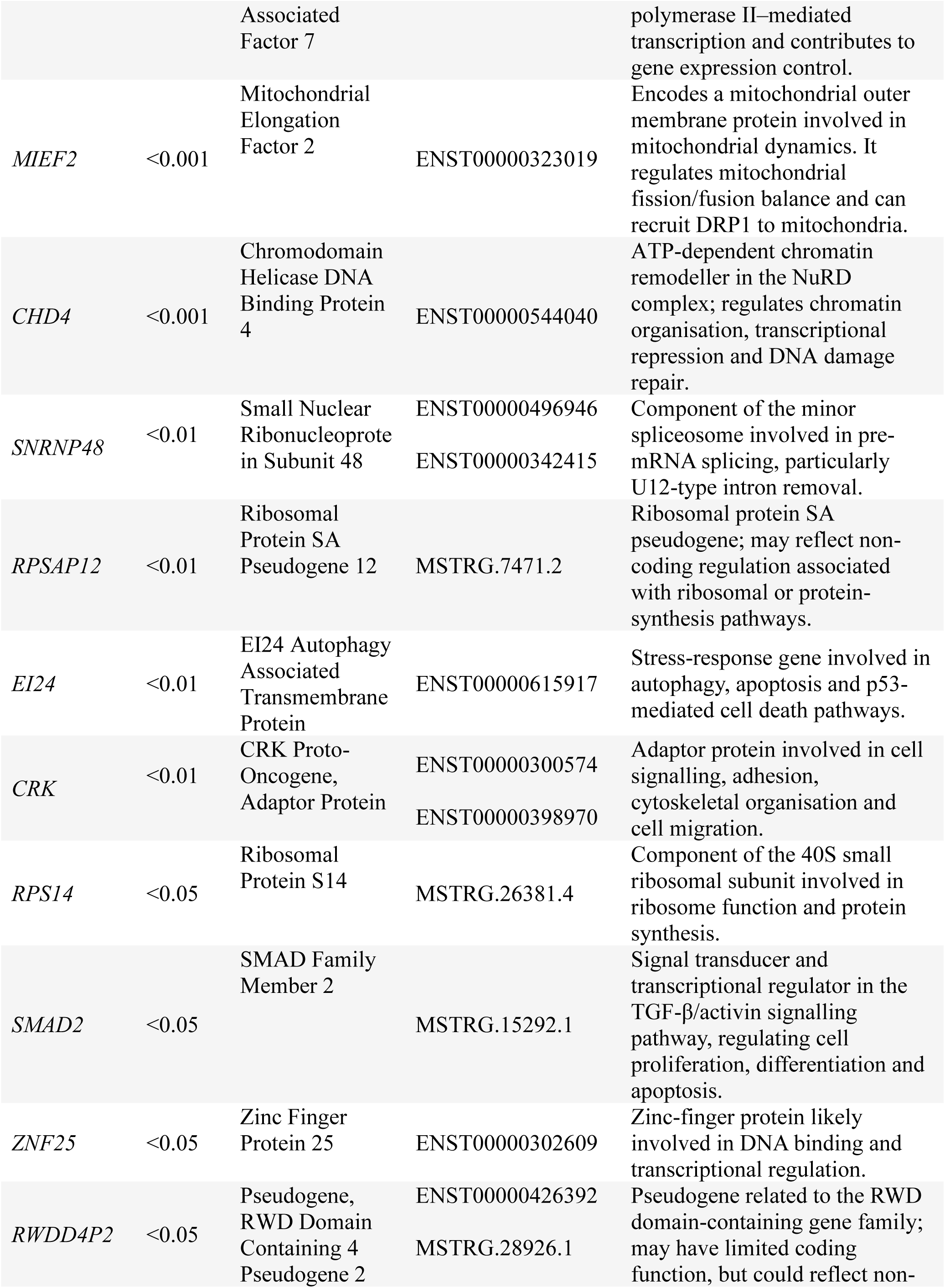
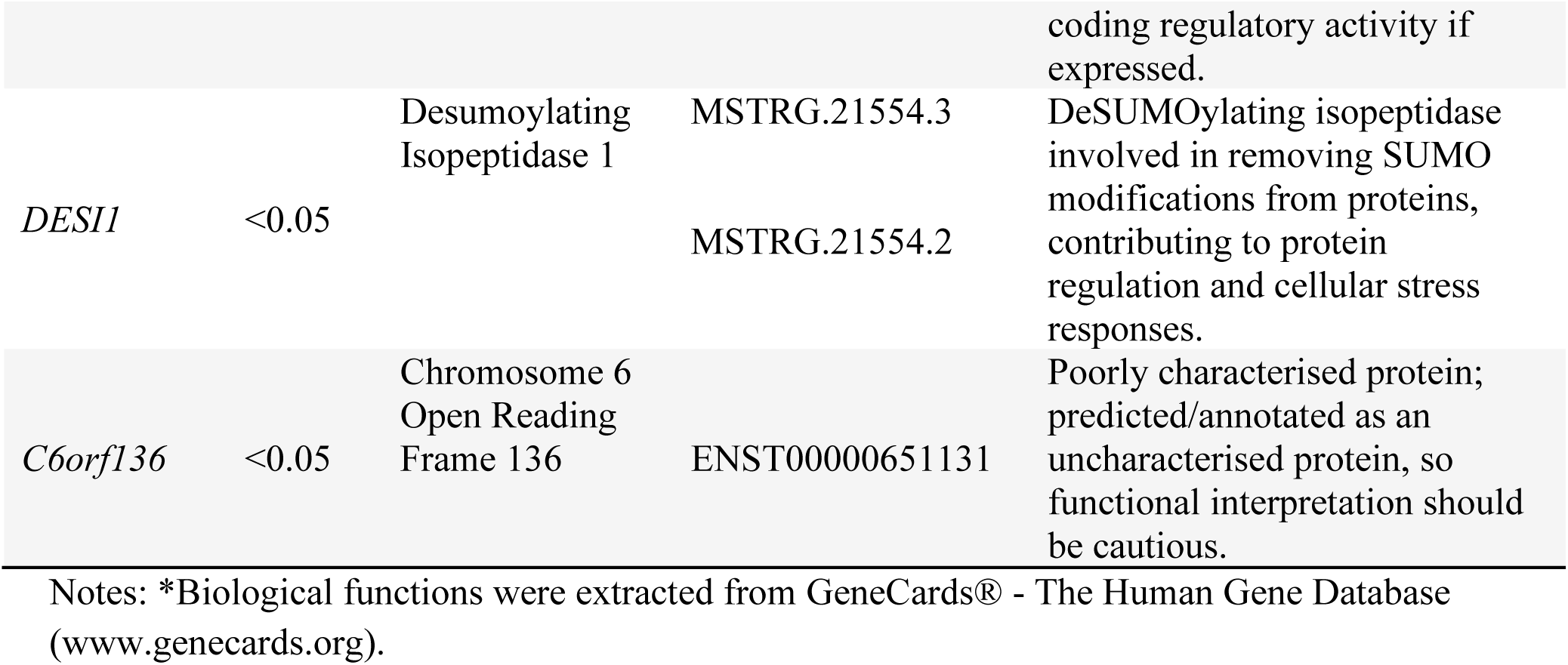
Genes with significant isoform switching and their functions. IDs with MSTRG are identified putative novel transcripts by StringTie.

The differential isoform-switching genes were associated with diverse biological functions (Table 2). These included genes involved in mitochondrial regulation and calcium handling (MIEF2 and CACNA1H); phosphorylation-dependent signalling, cytoskeletal organisation and cell adhesion (PPP1CC and CRK); RNA processing and non-coding or pseudogene-related regulation (SNRNP48); protein homeostasis and cellular stress responses (ERO1B); autophagy (EI24); and transcriptional and chromatin regulation (TAF7, CHD4, SMAD2 and ZNF25). To validate selected RNA-seq-identified isoform usage changes, we performed qRT-PCR analysis. However, because of limitations in identifying isoform-specific regions suitable for primer design, qRT-PCR validation was limited to selected isoforms of CACNA1H, SNRNP48, CRK and RWDD4P2 (Figure 2). qRT-PCR confirmed the direction of isoform usage changes for these selected genes, supporting the robustness of the RNA-seq findings. Together, these data suggest that isoform switching may represent an additional layer of molecular dysregulation in ME/CFS that is not fully captured by gene-level differential expression analysis alone.

**Figure 2.**
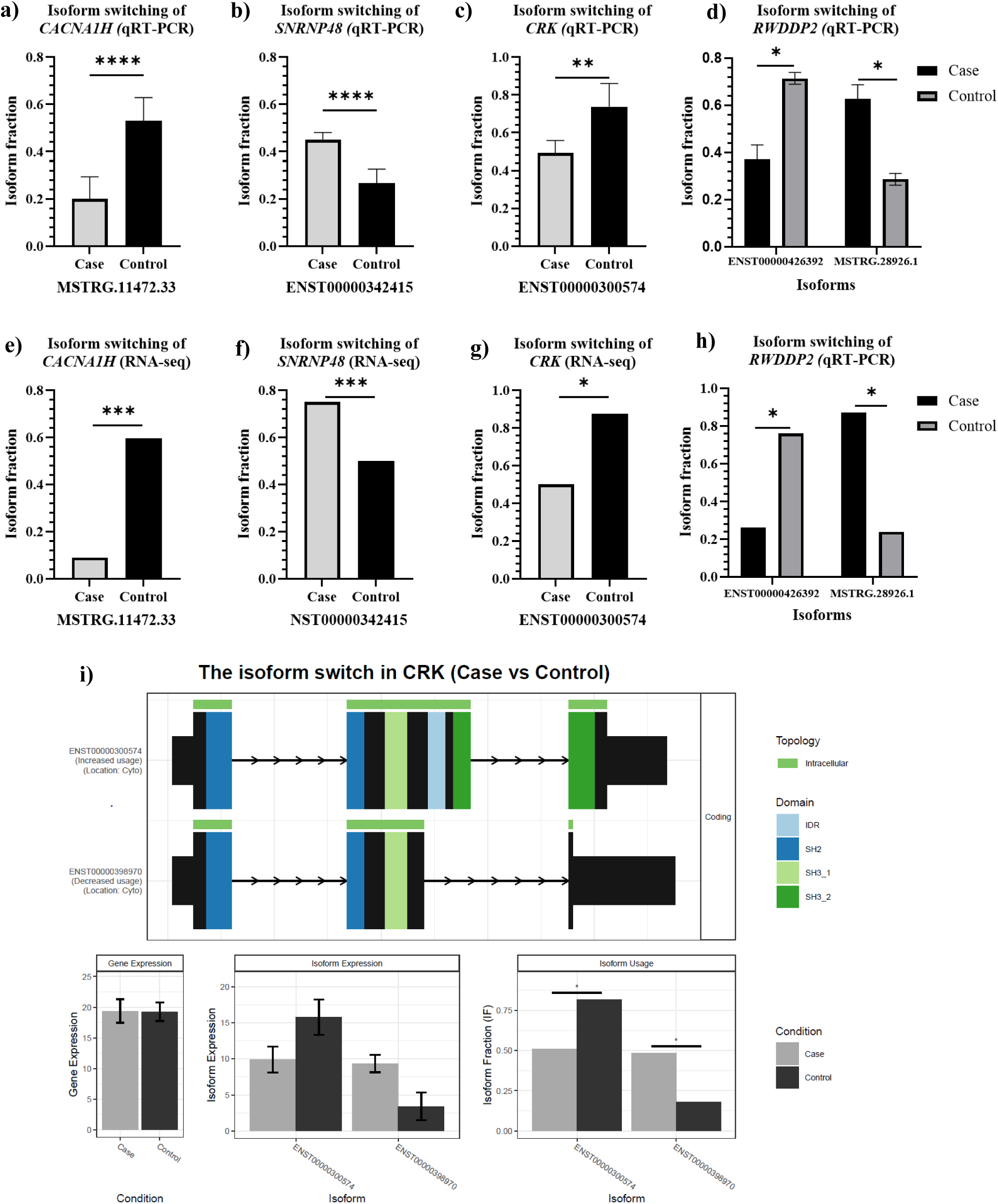
**Validation and structural characterisation of isoform switching in ME/CFS myotubes**. Isoform fractions of CACNA1H, SNRNP48, CRK and RWDD4P2 were assessed using qRT-PCR and compared with RNA-seq-derived isoform usage estimates. Isoform fractions of CACNA1H were determined by (a) qRT-PCR and (e) RNA-seq; SNRNP48 by (b) qRT-PCR and (f) RNA-seq; CRK by (c) qRT-PCR and (g) RNA-seq; and RWDD4P2 by (d) qRT-PCR and (h) RNA-seq. Panel (i) shows an example of predicted isoform switching and domain architecture for CRK between case and control myotubes. Full isoform domain plots for the 16 significant isoform-switching genes are provided in Supplementary Figure 2. Significant differences in qRT-PCR-derived isoform fraction were assessed using two-tailed unpaired Welch’s t-tests. Bars represent mean isoform fraction, and error bars indicate the standard error of the mean for qRT-PCR data.

## Pathway Enrichment Analysis

Pathway enrichment analysis identified multiple significantly enriched pathways in ME/CFS myotubes (Figure 3 and Supplementary File 3). The enriched pathways formed several functionally related clusters, including pathways associated with RNA metabolism, protein synthesis, protein processing, transport, metabolic regulation and cellular signalling. The pathway network also included pathways linked to cell signalling, immune-related processes and cellular communication, such as B cell receptor signalling pathway, cytosolic DNA-sensing pathway, GnRH secretion, insulin secretion, leukocyte transendothelial migration and vascular smooth muscle contraction. Pathways associated with neuronal or muscle-related processes were also observed, including motor proteins, axon guidance, spinocerebellar ataxia and taste transduction.

**Figure 3.**
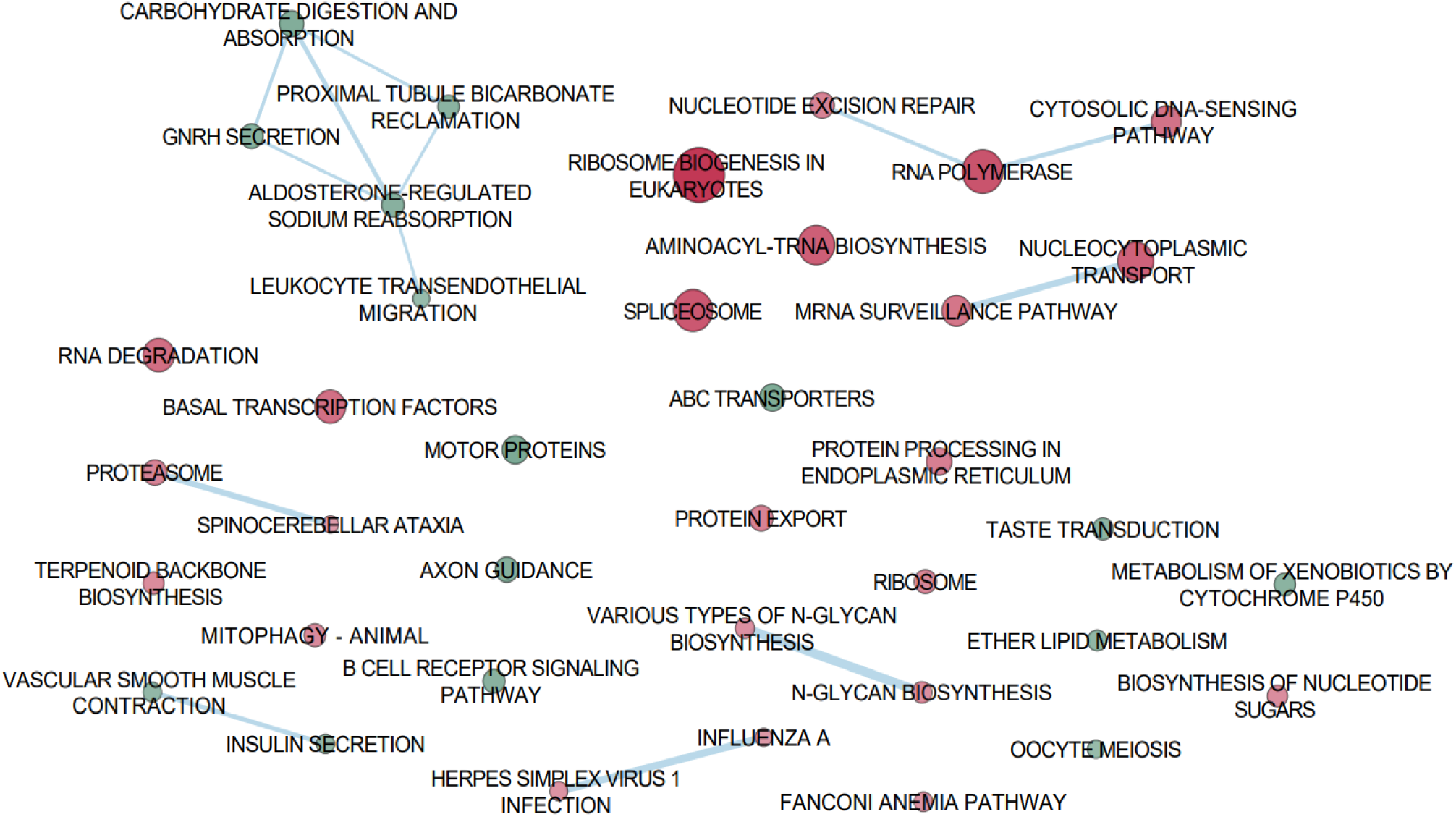
Enrichment Map of GSEA results in myotubes using KEGG pathways analysis, each node represents a significantly enriched pathway (q-value<0.05), node size reflects the absolute Normalized Enrichment Score (NES); green indicates positive NES (upregulated), and red indicates negative NES (downregulated), edge thickness corresponds to the number of overlapping genes between pathways. Additional information for GSEA is provided in the Supplementary File 3.

## Drug repurposing candidates for treatment of ME/CFS

Querying the LINCS2 database with the 100 upregulated and 100 downregulated most differentially expressed genes (all p < 0.01) identified several thousand compounds of interest (Supplementary File 4). Compounds with a weighted connectivity score false discovery rate (WTCS-FDR) below 0.05 were retained for further evaluation. These candidates were then reviewed in detail using a targeted literature search and filtered according to the predefined exclusion criteria. Fourteen compounds met the eligibility criteria, were approved for human use and were shortlisted as potential drug-repurposing candidates. The final candidate list is presented in Table 3, with supporting evidence for each compound summarised in Supplementary File 5.

**Table 3.** Selected drug candidates with repurposing potential for treatment of ME/CFS

| Drug | NCS | WTCS - FDR | Mechanism of action | Approved therapeutic indication |
| --- | --- | --- | --- | --- |
| Memantine | -1.86 | <0.0001 | NMDA receptor - Antagonist | Moderate to severe dementia of the Alzheimer |
| Riboflavin | -1.67 | <0.0001 | Vitamin B | Progressive keratoconus<br>Corneal ectasia |
| Pioglitazone | -1.51 | <0.0001 | Insulin sensitizer;<br>PPAR receptor agonist | X-linked adrenoleukodystrophy<br>Type 2 diabetes mellitus |
| Amantadine | -1.42 | <0.0001 | NMDA receptor - Antagonist | Influenza A virus infection<br>Parkinson's disease |
| Aniracetam | -1.41 | <0.0001 | Glutamate receptor ionotropic AMPA - Positive Allosteric Modulator | Senile dementia of the Alzheimer type |
| Benidipine | -1.39 | <0.0001 | Calcium channel blocker | Hypertension<br>Angina pectoris |
| Aprepitant | -1.36 | <0.0001 | Tachykinin antagonist | Postoperative nausea and vomiting |
| Bezafibrate | -1.29 | 0.000162 | Pan-PPAR ($\alpha/\delta/\gamma$) agonist | Hyperlipidaemia |
| Propranolol | -1.21 | <0.0001 | Adrenergic receptor antagonist | Angina pectoris<br>Hypertension<br>Hypertrophic subaortic stenosis |
| Fenofibrate | -1.19 | 0.000162 | PPARα agonist | Hypercholesterolemia or mixed dyslipidemia and hypertriglyceridemia |
| Acarbose | -1.18 | <0.0001 | Glucosidase inhibitor | Type 2 diabetes mellitus |
| Alfacalcidol | -1.12 | <0.0001 | Vitamin D receptor agonist | Rickets<br>Hyperparathyroidism |
| Nicotinamide | -1.11 | <0.0001 | NAD-dependent deacetylase sirtuin 1/2/3 - Inhibitor | Pellagra<br>Arteriosclerotic<br>Ophthalmology |
| Acyclovir | -1.11 | <0.0001 | DNA polymerase inhibitor | Cold sores<br>Genital herpes<br>Herpes zoster infections |

**NCS** = Normalised Connectivity Score. This value represents the strength of the connection between a drug-induced gene expression signature and the ME/CFS gene expression profile. A more negative NCS indicates a greater potential for the drug to reverse the disease signature. The shortlisted drugs are ranked by NCS values.
**WTCS-FDR** = Weighted connectivity Score-False discovery rate. The WTCS\_FDR represents the adjusted p-value (FDR) for a drug's connectivity score, estimating how likely the observed match/mismatch between a compound and a gene signature is due to chance.

## Discussion

Despite clinical trials evaluating pharmacological agents, nutraceuticals and exercise-based interventions, no treatment has been approved specifically for PEM in ME/CFS, reflecting the limited understanding of its underlying pathophysiology and the lack of validated therapeutic targets. In this study, we generated iPSC-derived myotubes as a participant-specific model for investigating ME/CFS biology. Under basal culture conditions, ME/CFS myotubes showed relatively modest differences in overall gene expression but more pronounced dysregulation at the transcript-isoform level. Pathway analysis identified 40 significantly altered KEGG pathways, while LINCS-based drug-signature mapping prioritised 14 compounds with potential for repurposing in ME/CFS.

## Transcriptomic and pathway enrichment landscape

Only a small number of genes met the genome-wide threshold for differential expression between ME/CFS and control myotubes, with three differentially expressed genes identified (ERVH48-1, CDX4 and SLC2A14). This limited differentially expressed genes profile suggests that disease-associated changes may not be strongly reflected in total gene abundance under basal culture conditions. This is consistent with the broader concept that ME/CFS-associated molecular abnormalities may be context-dependent and may become more apparent following physiological challenge, such as exercise or exertion [37, 38]. Among the differentially expressed genes , SLC2A14 encodes a member of the facilitated glucose transporter family [39], suggesting a possible link to altered substrate transport or metabolic regulation. CDX4 is a homeobox transcription factor involved in developmental gene regulation [40], while ERVH48-1 encodes an endogenous retrovirus-derived protein [41]. However, the functional relevance of these genes in mature skeletal muscle and ME/CFS remains unclear and requires further investigation.

In contrast to the gene expression findings, transcript isoform analysis identified 16 significant isoform-switching genes differentially regulated in ME/CFS, including CACNA1H, SNRNP48, CRK, RWDD4P2, PPP1CC, SMAD2, CHD4 and MIEF2. Alternative splicing and isoform switching are important mechanisms for expanding proteomic diversity and regulating skeletal muscle development, function and disease biology [42, 43]. Therefore, altered isoform usage may affect protein domains, regulatory motifs, subcellular localisation or transcript stability, even when total gene abundance is unchanged. CACNA1H is of particular interest because it encodes the CaV3.2 T-type calcium channel, which has been implicated in calcium signalling during human myoblast differentiation and fusion and in the maintenance of skeletal muscle integrity [44, 45]. More broadly, calcium-dependent signalling regulates skeletal muscle contraction, metabolic and mitochondrial adaptation, and cellular stress responses [46]. MIEF2 also has strong biological relevance to skeletal muscle function, as it encodes a mitochondrial outer membrane protein involved in mitochondrial morphology and fission, processes that are critical for skeletal muscle energy metabolism and exercise tolerance [47]. In addition, SMAD2 plays a key role in TGFβ signalling and skeletal muscle regulation. Yamasaki *et al*. (2026) reported that impaired SMAD2 ubiquitination increased SMAD2 protein levels and TGFβ responsiveness in skeletal muscle, leading to reduced muscle mass, smaller myofibre size and suppressed terminal differentiation of primary myoblasts [48]. The same study also reported enhanced fibrotic remodelling following muscle injury, supporting a role for SMAD2 regulation in both muscle homeostasis and tissue repair [48]. In this context, the SMAD2 isoform-switching event observed in ME/CFS iPSC-derived myotubes may reflect altered TGFβ/SMAD-related regulatory programmes with potential relevance to myogenic differentiation, myotube formation and muscle remodelling.

CHD4 is also of interest because of its established role in striated muscle identity and metabolic homeostasis. This gene plays an important role in the Chd4/NuRD chromatin-remodelling complex, which preserves cardiac and skeletal muscle identity by repressing inappropriate alternate lineage gene programmes [49]. In mouse skeletal muscle, CHD4 deletion caused smaller tibialis anterior muscles, features of myopathy, abnormal expression of cardiac-lineage genes, disrupted mitochondrial activity, and metabolic homeostasis in striated muscle [49].

Therefore, the CHD4 isoform-switching event observed in ME/CFS iPSC-derived myotubes may reflect altered epigenetic regulation of muscle identity and metabolic homeostasis, with potential relevance to skeletal muscle dysfunction in ME/CFS. CRK is also relevant to skeletal muscle function through its role in neuromuscular junction formation and maintenance. Klockner et al. (2022) demonstrated that CRK participates in a DOK7–CRK–RAC1 signalling cascade required for acetylcholine receptor clustering and neuromuscular junction function, with dysregulated CRK expression leading to impaired neuromuscular function [50]. Although our myotube model does not directly assess innervation, CRK isoform switching may point to altered cytoskeletal or adaptor-protein signalling pathways with potential relevance to neuromuscular organisation, muscle fatigue and post-exertional malaise.

The observation that isoform-level dysregulation was more prominent than gene-level differential expression suggests that ME/CFS myotubes may show subtle transcriptional changes at the total gene-expression level but clearer disease-associated changes at the transcript isoform level. This distinction is important because many transcriptomic studies focus primarily on differentially expressed genes and may overlook alternative splicing or isoform usage changes [24]. In ME/CFS, where biological abnormalities may be subtle, heterogeneous or context-dependent, isoform-level analysis may provide additional insight into disease mechanisms. These findings suggest that future studies should integrate gene-level, isoform-level and functional analyses to better understand disease-associated muscle biology.

Gene set enrichment analysis identified several coordinated pathway-level alterations with potential relevance to skeletal muscle function. Enrichment of motor-protein and vascular smooth muscle contraction pathways may reflect altered regulation of cytoskeletal, ion-handling and contractile processes rather than indicating a vascular smooth muscle phenotype. This interpretation is consistent with previous evidence of altered exercise-responsive proteins involved in muscle contraction and cytoskeletal organisation in ME/CFS, including myosin light chains, tropomyosin and tropomodulin, some of which were associated with post-exercise myalgia and fatigue [51]. Disturbed skeletal-muscle ion homeostasis has also been reported, with increased muscle sodium content before and after exercise and higher sodium levels associated with reduced handgrip strength [52].

Mitophagy was negatively enriched in ME/CFS myotubes, suggesting altered regulation of mitochondrial quality-control processes. Mitophagy is required for selective removal of dysfunctional mitochondria and maintenance of a healthy mitochondrial pool in skeletal muscle, particularly during adaptation to metabolic and contractile stress [53]. This finding is relevant to growing evidence of altered skeletal-muscle bioenergetics in ME/CFS. Participant-derived skeletal-muscle cells have shown abnormalities in mitochondrial substrate utilization [54], while skeletal-muscle biopsy and experimental studies have implicated disrupted mitochondrial respiratory-supercomplex formation and reduced respiratory function in ME/CFS [8]. Together, these pathway-level findings point to coordinated alterations in contractile regulation, ion homeostasis and mitochondrial quality control that may contribute to skeletal-muscle dysfunction in ME/CFS, although functional studies are required to establish their physiological consequences.

## Potential candidates for drug repurposing

An integrated computational analysis of transcriptomic signatures, LINCS2 drug-query results, and literature-based screening identified 14 potential drug candidates for future translational and clinical investigation. Although these candidates should not be interpreted as direct therapeutic recommendations, they suggest that the transcriptional state of ME/CFS myotubes may be pharmacologically modifiable through pathways related to mitochondrial bioenergetics, lipid and glucose metabolism, calcium/glutamate signalling, neuroimmune activation, and antiviral mechanisms.

A major theme emerging from the shortlisted compounds was the regulation of mitochondrial metabolism and cellular energy homeostasis. This is highly relevant to ME/CFS, as recent studies using skeletal muscle biopsies have reported evidence of mitochondrial dysfunction, including reduced mitochondrial respiratory capacity under hyperoxic conditions, decreased succinate dehydrogenase activity, and altered TCA-cycle-related metabolites, particularly following exercise-induced PEM [7, 55]. Other studies have also identified mitochondrial morphological abnormalities in ME/CFS and post-COVID-related fatigue syndromes, including altered cristae structure and subsarcolemmal mitochondrial changes [56, 57]. Hence, riboflavin and nicotinamide are particularly relevant because they support core mitochondrial redox biology. Riboflavin is the precursor of flavin mononucleotide and flavin adenine dinucleotide, which are required cofactors for multiple flavoproteins involved in oxidative phosphorylation and fatty acid oxidation [58]. High-dose riboflavin has shown clinical benefit in migraine prophylaxis, a condition in which mitochondrial energy metabolism has been implicated [59], supporting its broader relevance as a mitochondrial-supportive intervention. Nicotinamide, as a precursor in NAD+ metabolism, may influence redox balance, sirtuin activity, poly (ADP-ribose) polymerase (PARP)-dependent stress responses, and mitochondrial function [60]. However, the translational relevance of NAD+ precursor biology remains context-dependent, and not all human studies have demonstrated direct improvements in skeletal muscle function [61]. Therefore, riboflavin and nicotinamide may be best viewed as low-risk metabolic modulators that warrant further testing in patient-derived muscle models before translation into clinical trials for ME/CFS.

Several candidates converge on peroxisome proliferator-activated receptor signalling, particularly pioglitazone, fenofibrate and bezafibrate. PPAR pathways regulate lipid handling, fatty acid oxidation, mitochondrial biology, inflammation and insulin sensitivity, which are reported as relevant biological processes to skeletal muscle metabolism [62, 63]. Pioglitazone, a PPARγ agonist, has been shown in human skeletal muscle to increase adiponectin, stimulate AMPK signalling, and increase the expression of genes involved in mitochondrial function and fatty acid oxidation [64]. Fenofibrate, a PPARα agonist, and bezafibrate, a pan-PPAR agonist, have been linked experimentally to fatty acid oxidation and PGC-1α-associated mitochondrial biogenesis [63, 65, 66]. Bezafibrate has also been investigated in mitochondrial fatty acid oxidation disorders and mitochondrial myopathy, although clinical outcomes have been mixed [67, 68]. Therefore, PPAR-modulating drugs may be useful candidates for follow-up testing in ME/CFS myotubes, particularly under energetic stress conditions that better model PEM.

A second theme among the shortlisted candidates was modulation of glutamate and excitatory signalling, calcium homeostasis and neuroimmune-related pathways, including memantine, amantadine, aprepitant and benidipine. These drugs are primarily used or investigated in neurological, inflammatory or cardiovascular contexts, reflecting overlap between these clinical pathways and stress-response mechanisms relevant to skeletal muscle biology. Calcium-dependent signalling is central to excitation–contraction coupling, mitochondrial activation, oxidative metabolism and fatigue-related changes in skeletal muscle [69]; therefore, candidates affecting excitatory or calcium-associated signalling may be relevant for follow-up testing in ME/CFS myotubes under energetic or contraction-like stress conditions.

Memantine and amantadine are clinically used adamantane derivatives with NMDA receptor antagonist properties [70–72]. Memantine is approved for moderate-to-severe Alzheimer’s disease and is used to reduce pathological glutamatergic signalling and excitotoxicity [73]. Amantadine has dopaminergic and NMDA-modulating actions and has been investigated for fatigue in neurological conditions, particularly multiple sclerosis, although controlled evidence remains mixed [62, 74]. More recently, an open-label randomised study reported improvement in post-COVID fatigue after short-term amantadine treatment, although placebo-controlled studies are still needed [75].

Aprepitant and benidipine suggest additional mechanisms involving neuroimmune signalling and calcium-dependent stress responses. Aprepitant is a neurokinin-1 receptor antagonist that blocks substance P signalling [76]. Substance P has been implicated in pain transmission, inflammation, neurogenic inflammation and immune-cell activation, processes that may be relevant to ME/CFS symptoms such as pain, sensory hypersensitivity and post-exertional symptom exacerbation [77, 78]. Some studies have reported that aprepitant can reduce inflammatory signalling, including JNK, p38/MAPK and NF-κB pathways, and may attenuate oxidative stress in immune-cell models [79, 80]. Benidipine, a calcium channel blocker, may be relevant because calcium handling is central to skeletal muscle excitation-contraction coupling, mitochondrial activation and fatigue biology [69]. Benidipine has been reported to reduce oxidative stress in clinical and experimental settings, including in polymorphonuclear cells from patients with essential hypertension [81], but direct evidence in ME/CFS or skeletal muscle fatigue models remains limited. Therefore, these candidates can be prioritised as mechanistic probes for follow-up testing in ME/CFS myotubes before conclusions can be drawn regarding their clinical translation.

## Limitations and future directions

This study has several limitations. First, the relatively small number of independent donor-derived lines limited statistical power, particularly at the individual-gene and isoform levels, and both false-positive and false-negative findings remain possible. The all-female cohort also limits generalisability to males with ME/CFS. These findings should therefore be considered hypothesis-generating and require replication in larger, more diverse independent cohorts. Second, each donor was represented by a single iPSC clone, and each line underwent a single terminal myotube differentiation experiment, limiting assessment of clone- and differentiation-related variability. Directed myogenic differentiation may also introduce line- and batch-dependent variation in differentiation efficiency. Future studies should incorporate replicate clones and independent differentiations, together with quantitative assessment of myotube differentiation and maturity.

Finally, transcriptomic profiling and drug-repurposing analyses were performed under basal culture conditions, which may not capture abnormalities that emerge following exertion-like stress. The computationally prioritised compounds therefore require direct experimental validation. Future studies should test these candidates in ME/CFS-derived myotubes under controlled stress, such as electrical pulse stimulation, using functional readouts including mitochondrial respiration, metabolic responses, calcium handling and transcriptomic reversal. Such studies will help determine whether the identified molecular alterations and candidate compounds are relevant to disease-associated muscle dysfunction and PEM.

## Conclusion

This study establishes patient-derived iPSC myotubes as a model for investigating skeletal muscle-associated molecular alterations in ME/CFS. Although gene-level differences were modest, complementary isoform-switching and pathway analyses identified broader alterations involving calcium signalling, mitochondrial quality control, metabolism and muscle-related processes, while computational drug repurposing prioritised candidate compounds for further testing. These findings provide a foundation for future functional studies under exertion-like stress and experimental validation of potential therapeutic candidates.

## Acknowledgments

The authors sincerely thank all individuals with ME/CFS and healthy control participants who generously donated their time and biological samples to support this research. Their contributions have been essential in advancing our understanding of ME/CFS and enabling the development of patient-specific disease models. We are especially grateful to the Geelong ME/CFS Support Group and Emerge Australia for their valuable assistance with participant recruitment for this project, this work would not have been possible without their support.

## Declarations

### Ethics approval and consent to participate

Ethics approval for this study was obtained from the Barwon Health Human Research Ethics Committee (20/202) and the Deakin University Human Research Ethics Committee (2021-165), Australia. The study was conducted in accordance with the ethical requirements approved by these Human Research Ethics Committees and the Australian National Statement on Ethical Conduct in Human Research. Written informed consent was obtained from all participants before participation.

## Consent for publication

Not applicable.

## Data and code availability

Due to ethical and privacy restrictions, the data and code that support the findings of this study are available upon reasonable request from the corresponding author.

## Author Contributions

Conceptualisation: KW, MB, CB, JHK, DN, BP, TT Methodology:

Recruitment JB, JS

Lab-based experiments BP, ZL, BS, ME, CS, DN, CF Data analysis TT, DN, BP, KW

Writing: DN

Review and editing: All authors

Funding acquisition: KW, MB, JHK, CB, BP

## Lead contact

Requests for further information and resources should be directed to and will be fulfilled by the lead contact, Ken Walder.

## KEY RESOURCES TABLE

| REAGENT or RESOURCE | SOURCE | IDENTIFIER |
| --- | --- | --- |
| **Antibodies** |  |  |
| Rabbit anti-PAX6 | BioLegend | Cat# 901301; RRID: AB_2565003 |
| Mouse anti-Nestin | Abcam | Cat# ab22035; RRID: AB_446723 |
| Rabbit anti-Brachyury (T) | Thermo Fisher Scientific | Cat# PA5-114917; RRID: AB_2899553 |
| Mouse anti-NCAM (CD56) | STEMCELL Technologies | Cat# 60021PE; RRID: AB_2938621 |
| Mouse anti-AFP | Millipore | Cat# ST1673-100UG; RRID: AB_10693988 |
| Rabbit anti-SOX17 | Abcam | Cat# ab224637; RRID: AB_2801385 |
| Rabbit monoclonal anti-OCT4 | Abcam | Cat# ab181557; RRID: AB_2687916 |
| Mouse anti-TRA-1-60 | Applied Stemcell | Cat# ASA-0160; RRID: AB_2827667 |
| Mouse anti-Myogenin | Sigma-Aldrich | Cat# M5815; RRID: AB_477218 |
| Rabbit polyclonal anti-MyoD | Thermo Fisher Scientific | Cat# PA5-23078; RRID: AB_11152869 |
| Mouse monoclonal anti-myosin heavy chain (MyHC) | Abcam | Cat# B37484; AB_2921304 |
| Rabbit polyclonal anti-titin | Invitrogen | Cat# PA5-52379; RRID: AB_2648492 |
| Goat anti-Mouse IgG (H+L), Alexa Fluor 488 (secondary) | Invitrogen | Cat# A-11029; RRID: AB_2534088 |
| Donkey anti-Rabbit IgG (H+L), Alexa Fluor 594 (secondary) | Invitrogen | Cat# A-21207; RRID: AB_14163 |
| Goat anti-Mouse IgG (H+L), Alexa Fluor 647 (secondary) | Thermo Fisher Scientific | Cat# A32728; RRID: AB_2633277 |
| Goat anti-Chicken IgY (H+L), Alexa Fluor 488 (secondary) | Abcam | Cat# ab150169; RRID: AB_2636803 |
| Mouse monoclonal anti-SSEA-4 (IgG3), PE/Cy7 (flow cytometry) | BioLegend | Cat# 330419; RRID: AB_2629630 |
| Mouse IgG1 $\kappa$ Isotype control, PE/Cy7 | BioLegend | Cat# 400125; RRID: AB_2861433 |
| Mouse monoclonal anti-EpCAM (CD326) (IgG2b), BV421 | BioLegend | Cat# 324219; RRID: AB_11124342 |
| Mouse IgG2b Isotype control, BV421 | BioLegend | Cat# 400341; RRID: AB_10898160 |
| Mouse monoclonal anti-CD9 (IgG1), FITC | BD Biosciences | Cat#555371; RRID: AB_395773 |
| Mouse IgG1 Isotype control, FITC | BD Biosciences | Cat#555748; RRID: AB_396090 |
| **Bacterial and virus strains** |  |  |
| N/A | N/A | N/A |
| **Biological samples** |  |  |
| Blood from study participants (for PBMCs) | This paper | N/A |
| Human PBMCs from study participants (for iPSC generation) | This paper | N/A |
| **Chemicals, peptides, and recombinant proteins** |  |  |
| StemSpan SFEM II | STEMCELL Technologies | Cat#09605 |
| Lymphoprep | STEMCELL Technologies | Cat# 07811 |
| BD Horizon™ Fixable Viability Stain 570 | Becton Dickinson Science | Cat#564995 |
| Triton X-100 | Sigma-Aldrich | Cat#T-9284 |
| PBS | Gibco | Cat#10010023 |
| FBS | Gibco | Cat#A5670701 |
| 0.5M EDTA | Invitrogen | Cat#15575038 |
| NucBlue® (Hoechst 33342) | Invitrogen | Cat# R37605 |
| Qiagen RNeasy kit | Qiagen | Cat# 74104 |
| Maxima H Minus first strand cDNA synthesis kit | Thermo Fisher Scientific | Cat#K1651 |
| 16% Paraformaldehyde | ProSciTech | Cat#C004 |
| Bovine serum albumin solution | Sigma-Aldrich | Cat#A8412 |
| STEMdiff™ Trilineage Differentiation Kit | STEMCELL Technologies | Cat# 05230 |
| StemSpan Erythroid Expansion Supplement (100X) | STEMCELL Technologies | Cat#02692 |
| StemSpan™ SFEM II medium | STEMCELL Technologies | Cat# 09655 |
| Geltrex LDEV-Free Reduced Growth Factor Basement Membrane Matrix | Gibco | Cat#A1413302 |
| mTeSR Plus Kit | STEMCELL Technologies | Cat#100-0276 |
| Y-27632 (ROCK inhibitor) | STEMCELL Technologies | Cat#100-0483 |
| ReLeSR | STEMCELL Technologies | Cat#100-0151 |
| DMEM/F-12, GlutaMAX supplement | Gibco | Cat#10565018 |
| TrypLE™ Select Enzyme | Gibco | Cat#12563029 |
| Collagenase Type IV (1 mg/mL) | STEMCELL Technologies | Cat#07909 |
| D-PBS (without Ca<sup>2+</sup>/Mg<sup>2+</sup>) | Gibco | Cat#14190144 |
| MyoCult-SF Expansion Supplement (Human) | STEMCELL Technologies | Cat#05980 |
| STEMdiff™ Myogenic Progenitor Supplement Kit | STEMCELL Technologies | Cat#100-0151 |
| MyoCult™ Differentiation Kit (Human) |  | Cat#05965 |
| DMEM with 1000 mg/L D-Glucose | STEMCELL Technologies | Cat#36253 |
| CryoStor CS10 | STEMCELL Technologies | Cat#07930 |
| StemPro Accutase Cell Dissociation Reagent | Gibco | Cat#A1110501 |
| Corning Matrigel hESC-Qualified Matrix | Corning | Cat#354277 |
| **Critical commercial assays** |  |  |
| MycoAlert® Mycoplasma Detection Kit | Lonza Bioscience | Cat#LT07-318 |
| MycoAlert Assay Control Set | Lonza Bioscience | Cat#LT07-518 |
| The Quant-iT™ OliGreen® ssDNA Assay Kit | Invitrogen | Cat# O11492 |
| **Deposited data** |  |  |
| RNA-seq dataset from this study | This paper | Available on request |
| **Experimental models: Cell lines** |  |  |
| Human iPSC lines (generated from PBMCs) | This paper | N/A |
| Human iPSC-derived Myogenic Progenitors (MPs) | This paper | N/A |
| **Experimental models: Organisms/strains** |  |  |
| N/A | N/A | N/A |
| **Oligonucleotides** |  |  |
| Primers for 13 genes, see Supplementary Table 2 | This paper | N/A |
| **Recombinant DNA** |  |  |
| Epi5 Epitomal iPSC reprogramming vector (includes Epi5 Reprogramming Vector and Epi5 p53 & EBNA Vector) | Invitrogen/Thermo Fisher Scientific | Cat#A15960 |
| **Software and algorithms** |  |  |
| signatureSearch | Duan et al. [18] | https://doi.org/10.1093/nar/gkaa878 |
| fastp | Chen et al. [21] | https://doi.org/10.1093/bioinformatics/bty560 |
| HISAT2 | Kim et al. [22] | https://doi.org/10.1038/s41587-019-0201-4 |
| featureCounts | Liao et al. [23] | https://doi.org/10.1093/bioinformatics/btt656 |
| DESeq2 package in R | Love et al. [24] | https://doi.org/10.1186/s13059-014-0550-8 |
| sva package in R | Leek et al. [25] | https://doi.org/10.1093/bioinformatics/bts034 |
| clusterProfiler package in R | Yu et al. [26] | https://doi.org/10.1089/omi.2011.0118 |
| Cytoscape platform | Shannon et al. [27] | https://doi.org/10.1101/gr.1239303 |
| StringTie | Pertea et al. [28] | https://doi.org/10.1093/nar/gkt006 |
| CPAT | Mistry et al. [29] | https://doi.org/10.1038/s41587-021-01156-3 |
| Pfam-A database | Mistry et al. [30] | https://doi.org/10.1093/nar/gkaa913 |
| SignalP 6.0 | Teufel et al. [31] | https://doi.org/10.1038/s41587-021-01156-3 |
| DeepLoc2 | Thumuluri et al. [32] | https://doi.org/10.1093/nar/gkac278 |
| DeepTMHMM | Hallgren et al. [33] | https://doi.org/10.1101/2022.04.08.487609 |
| IUPred3 | Erdős et al. [34] | https://doi.org/10.1093/nar/gkab408 |
| IsoformSwitchAnalyzeR | Vitting-Seerup and Sandelin, [35] | https://doi.org/10.1093/bioinformatics/btz247 |
| SatuRn | Gilis et al. [36] | https://doi.org/10.12688/f1000research.51749.2 |
| GraphPad Prism version 10.3.1 | GraphPad | Prism 10.3.0 |
| **Other** |  |  |

---

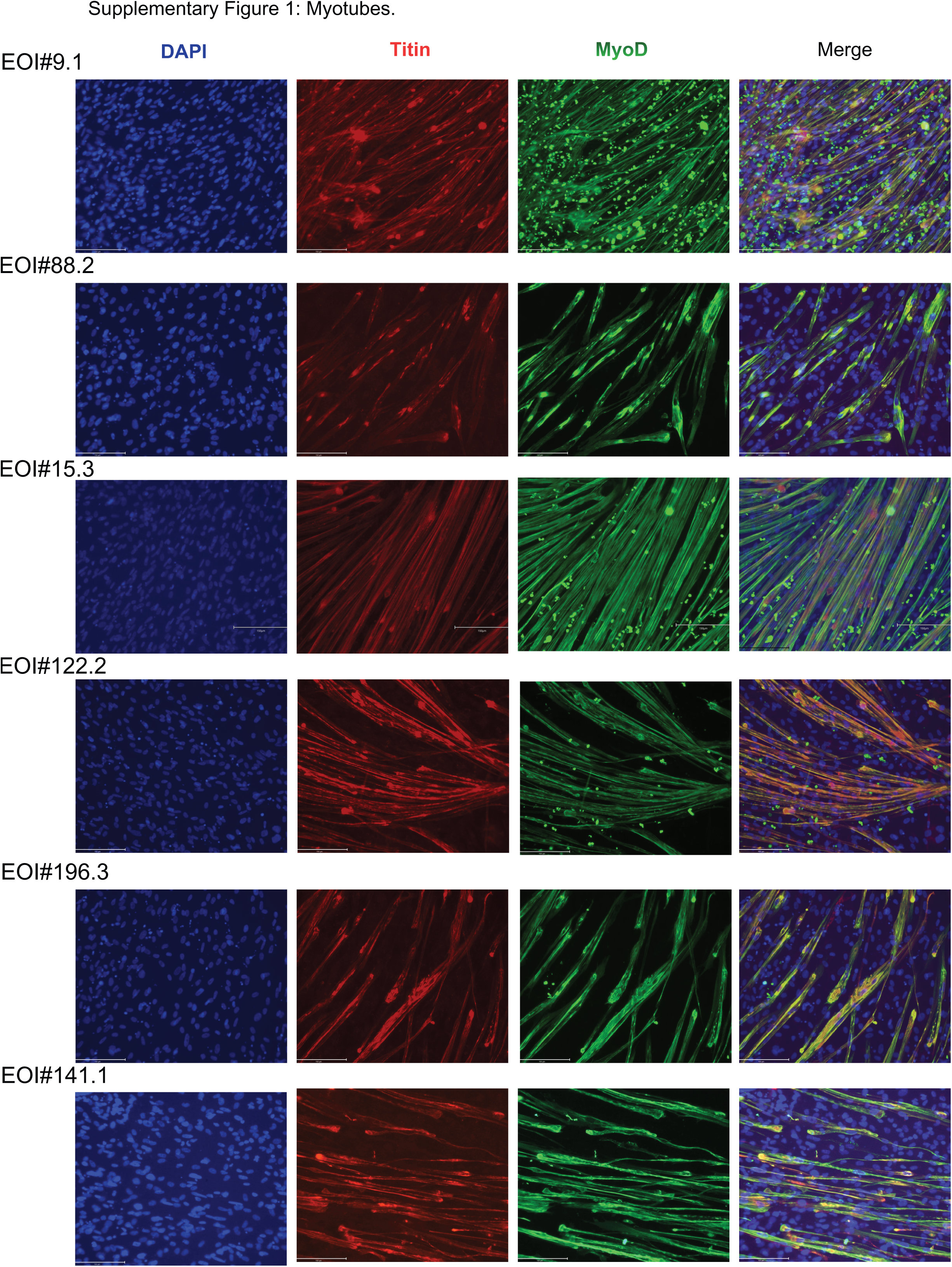
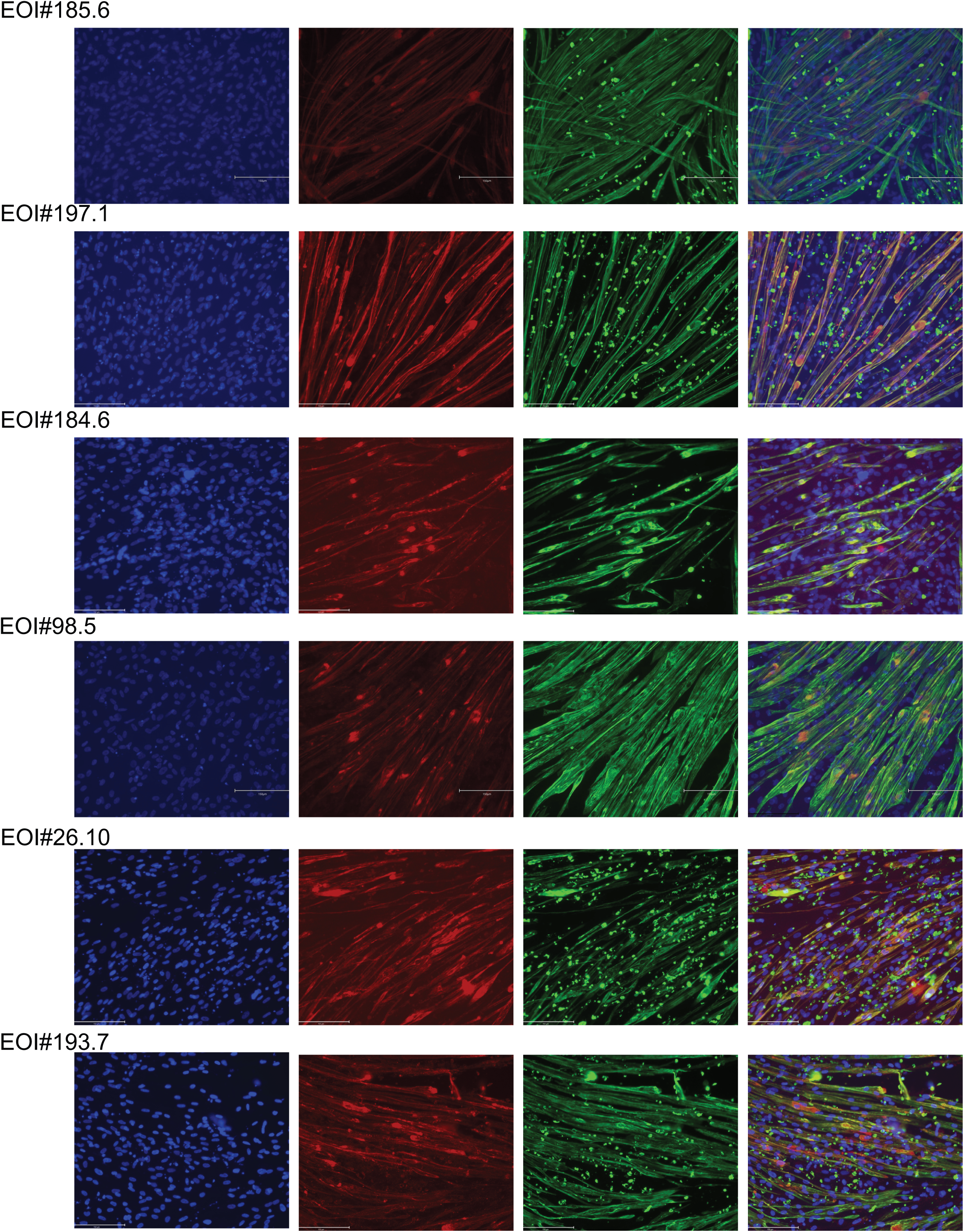
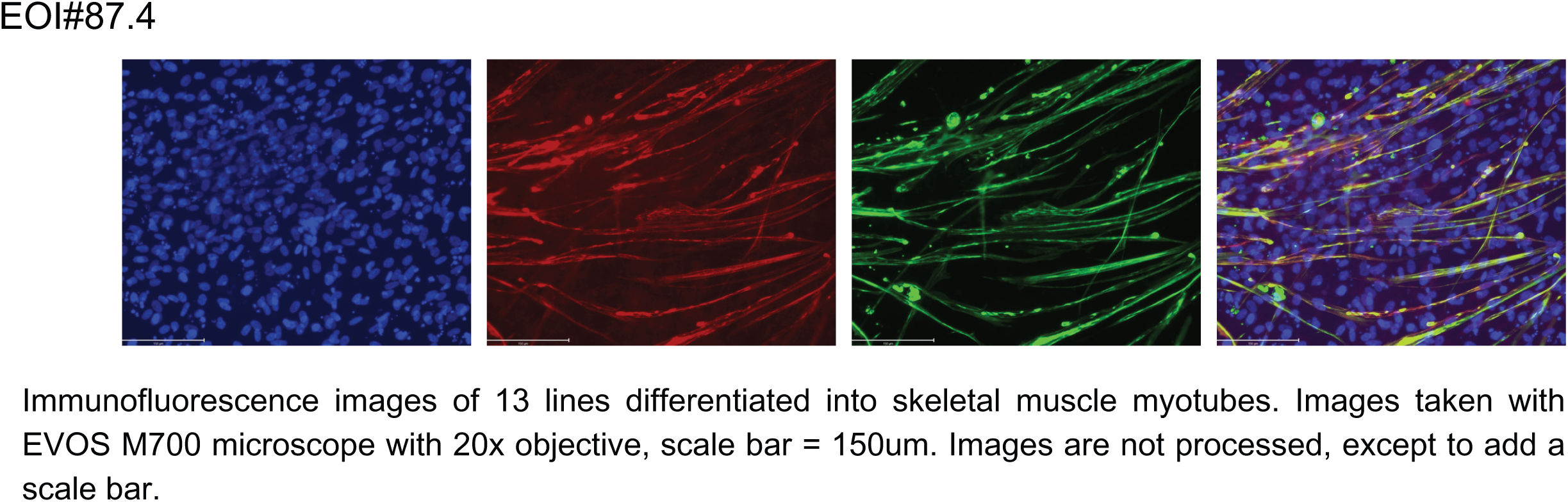

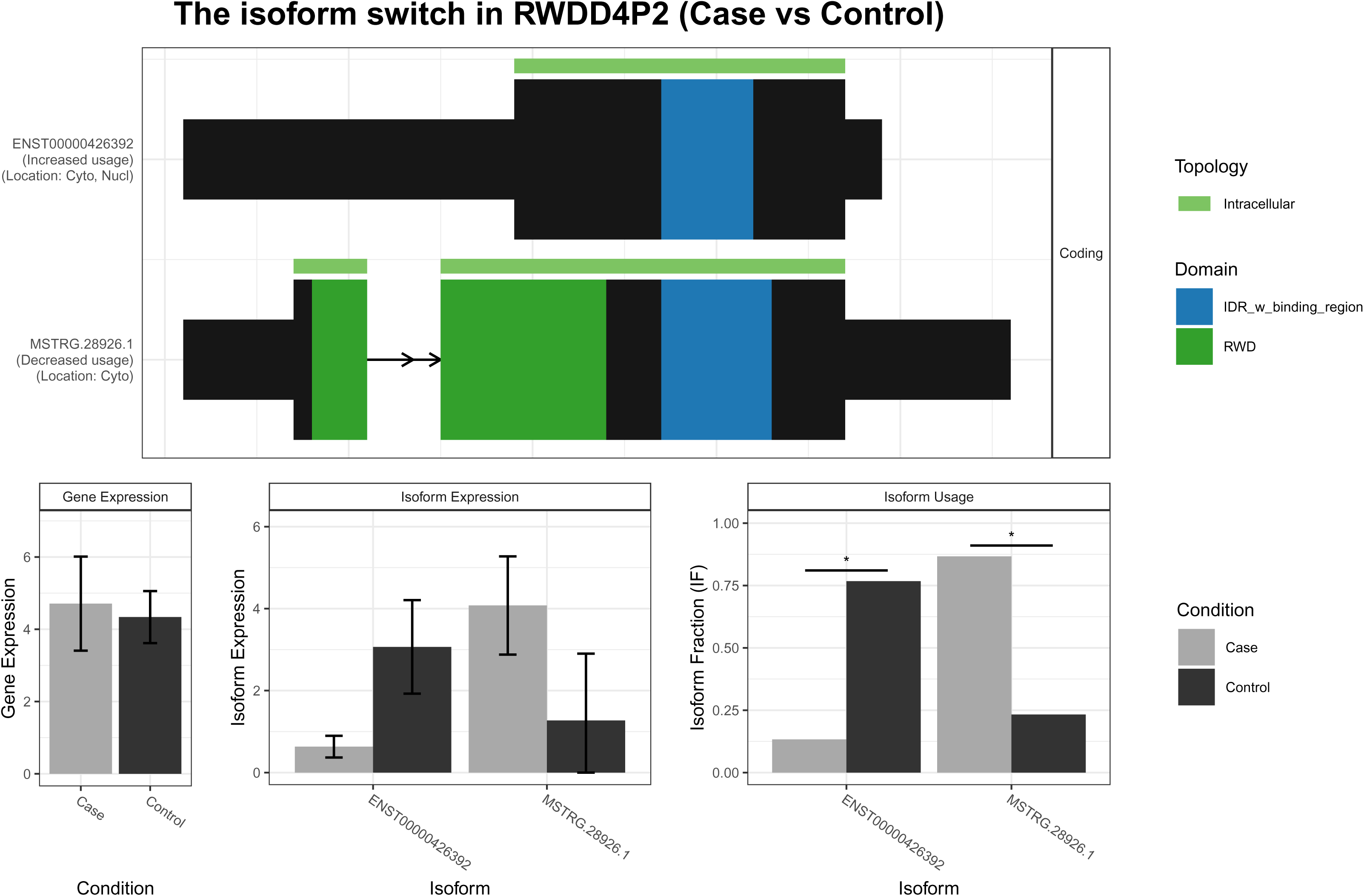

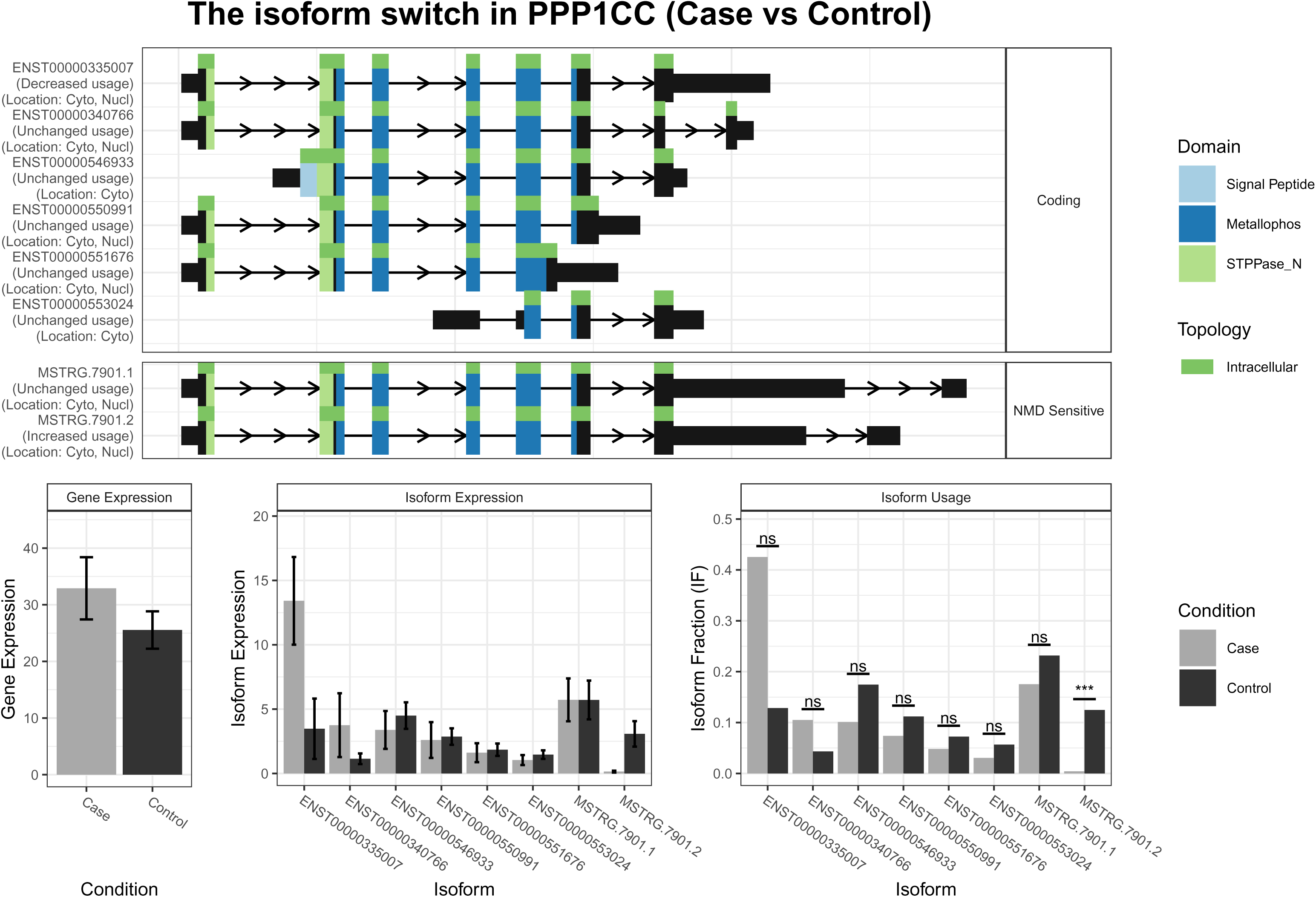

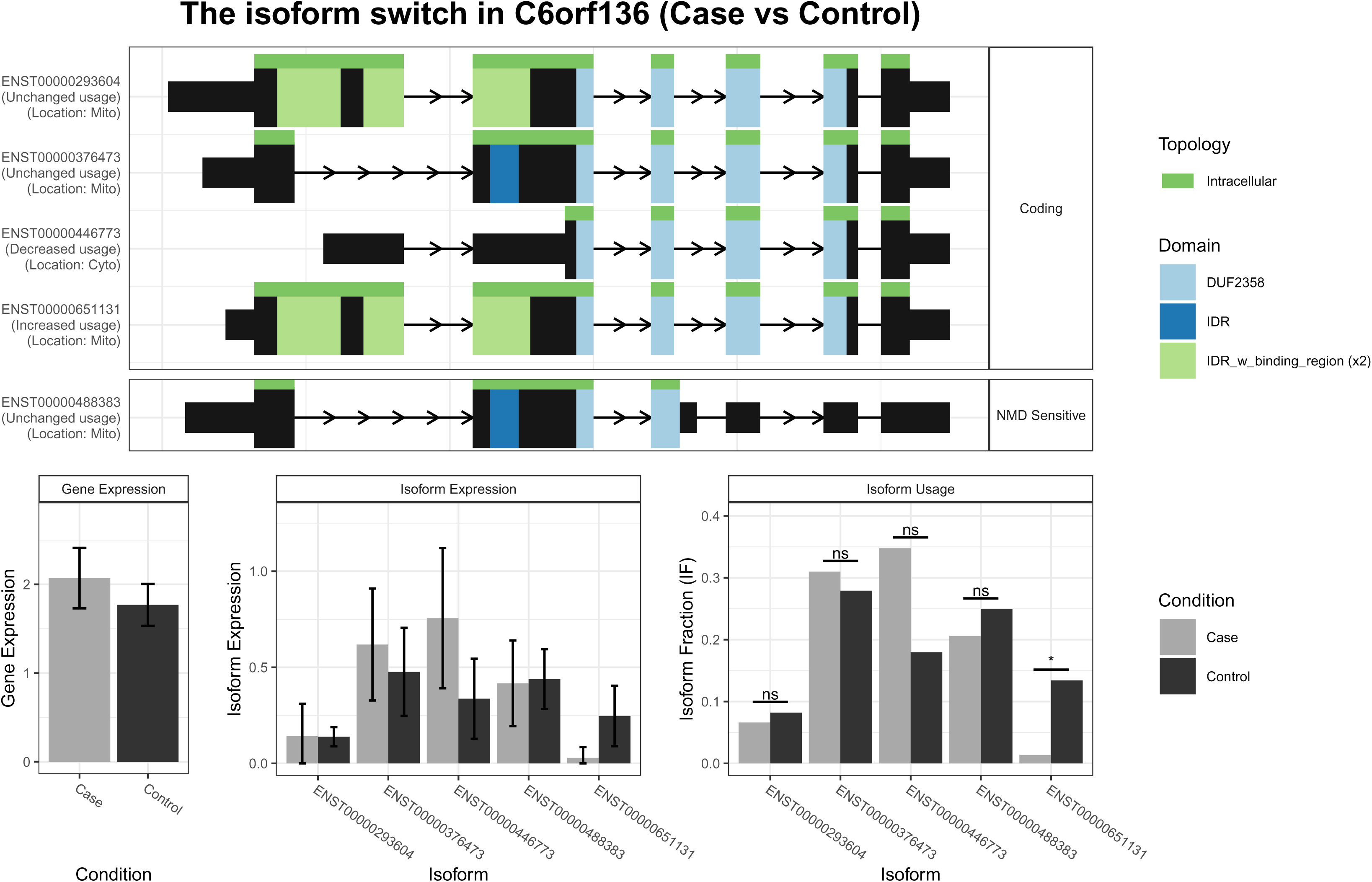

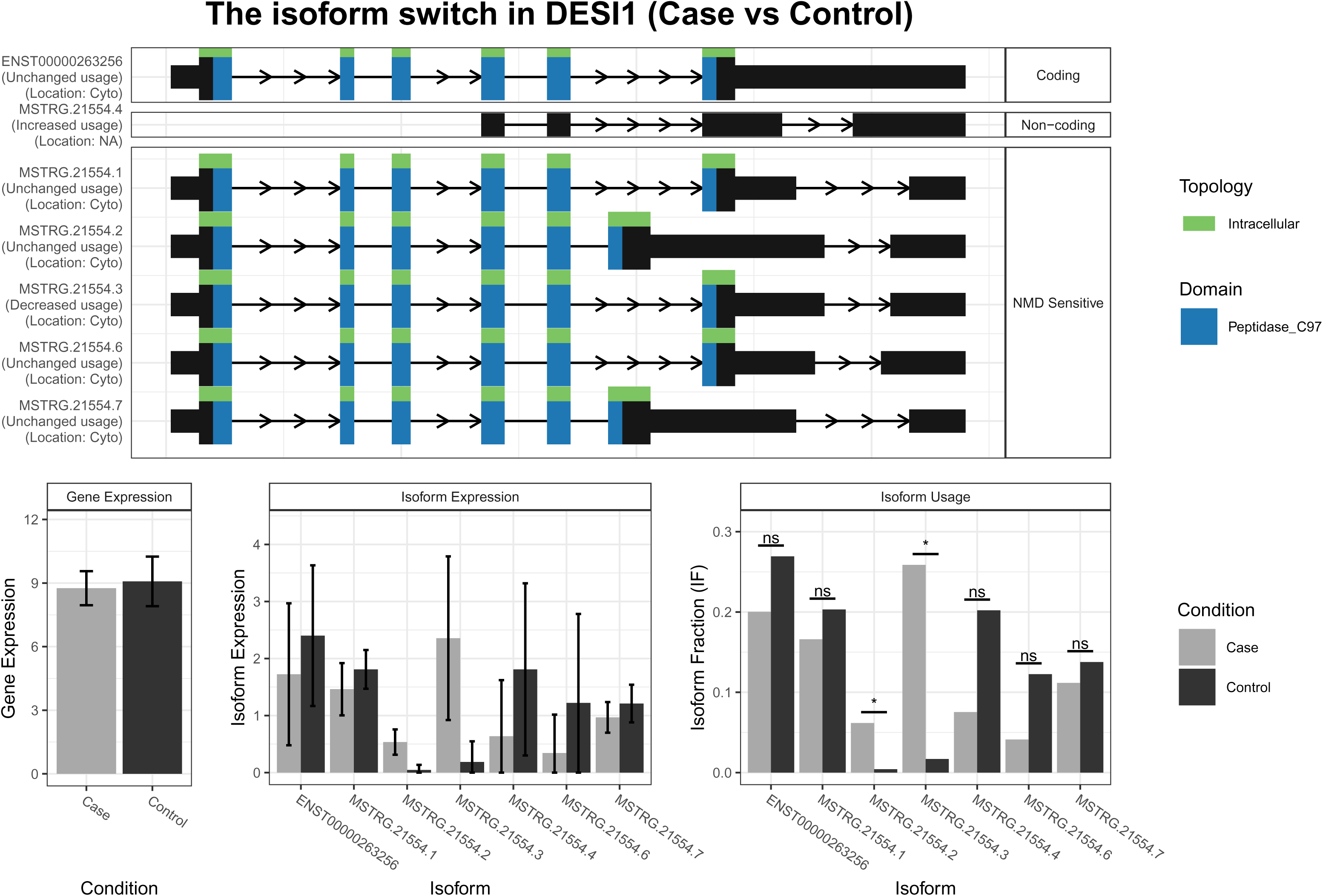

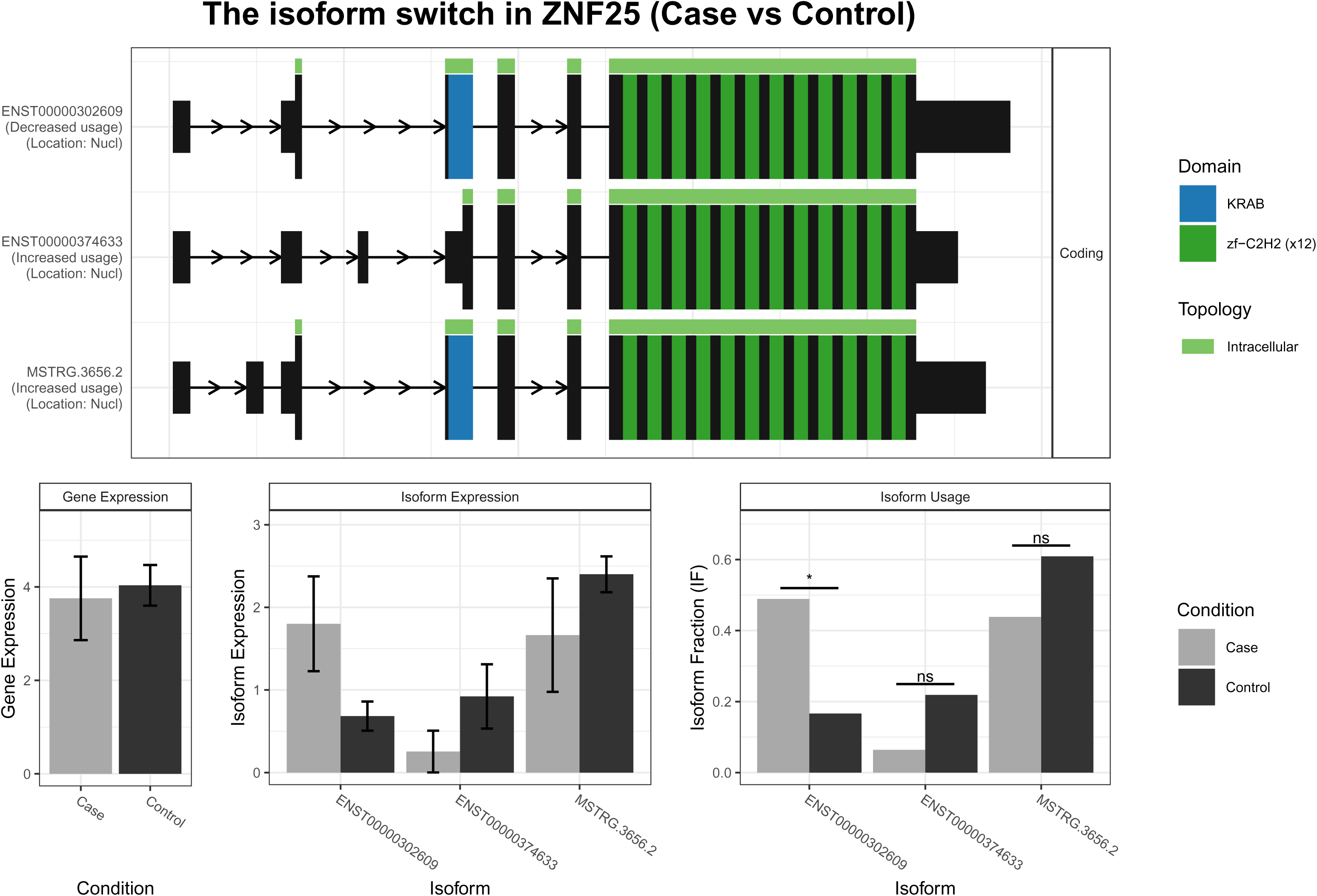

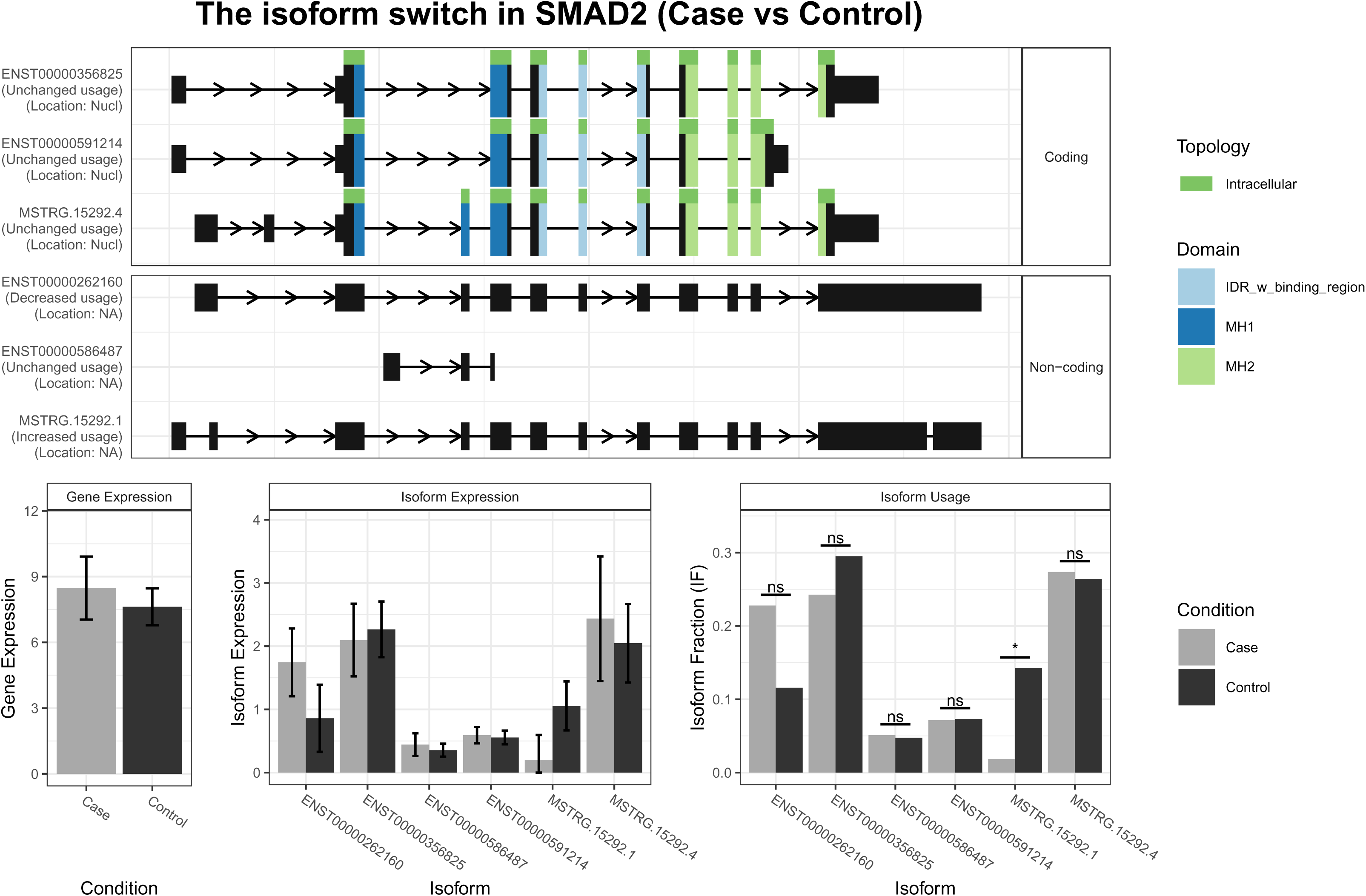

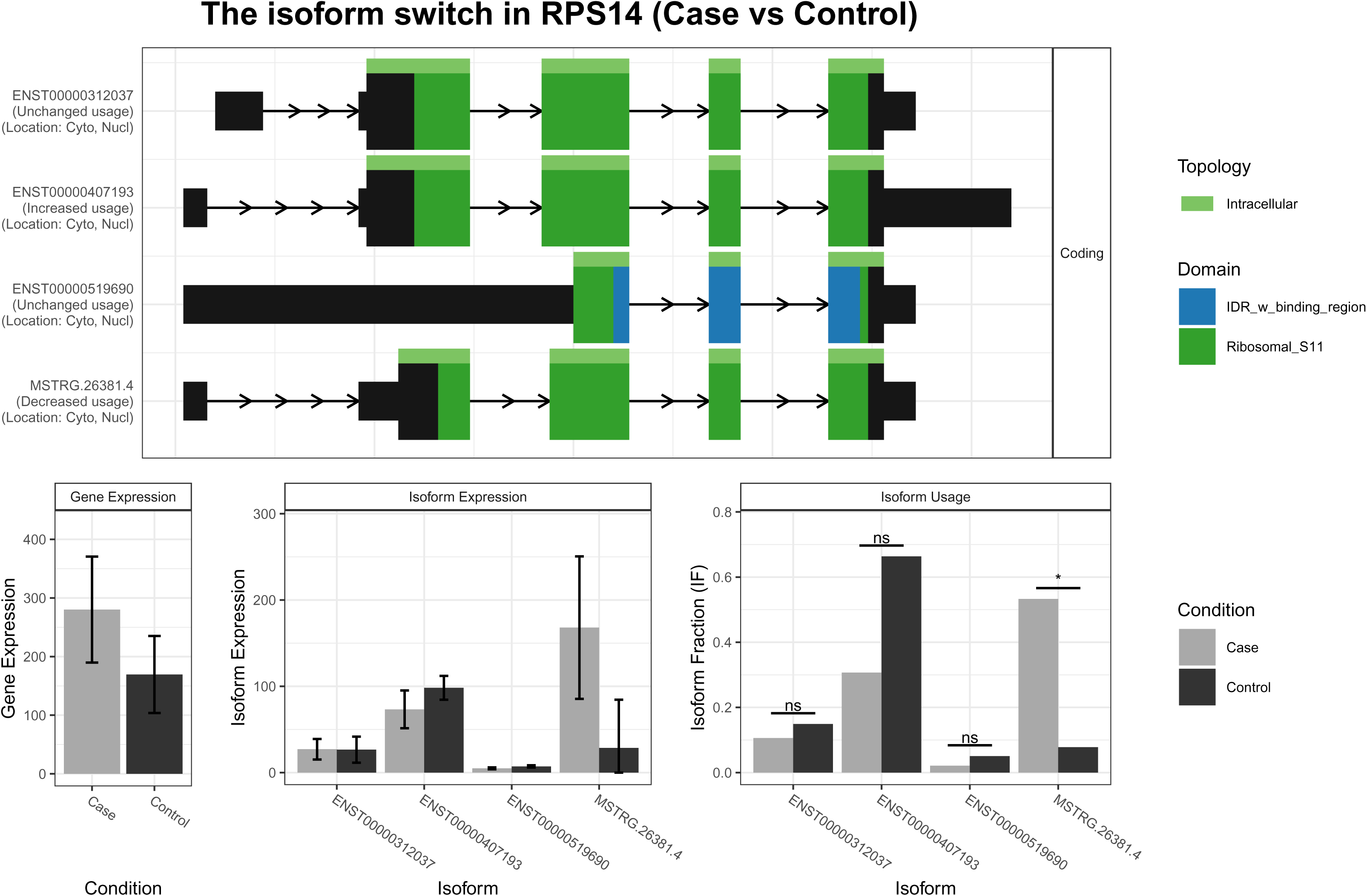

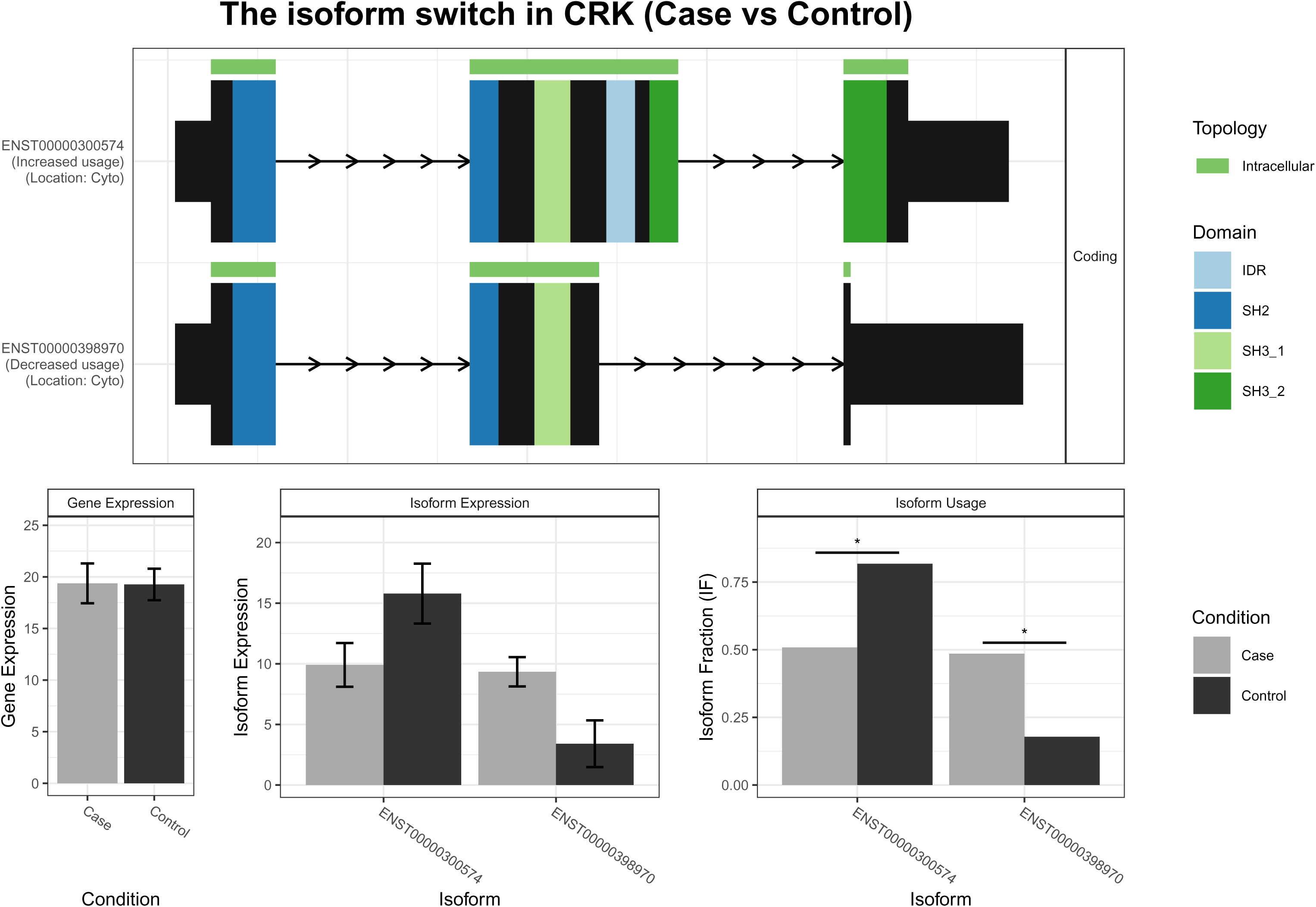

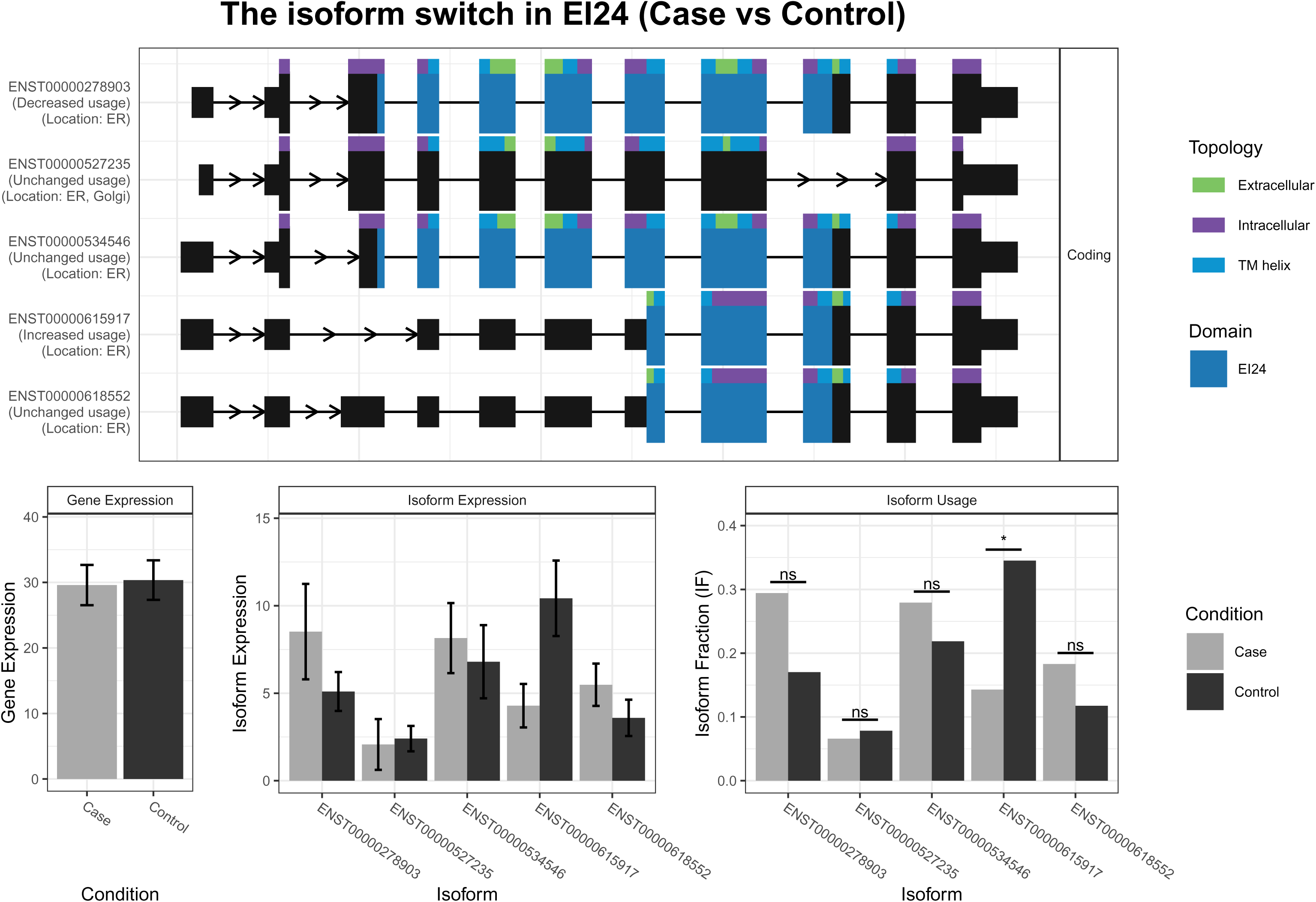

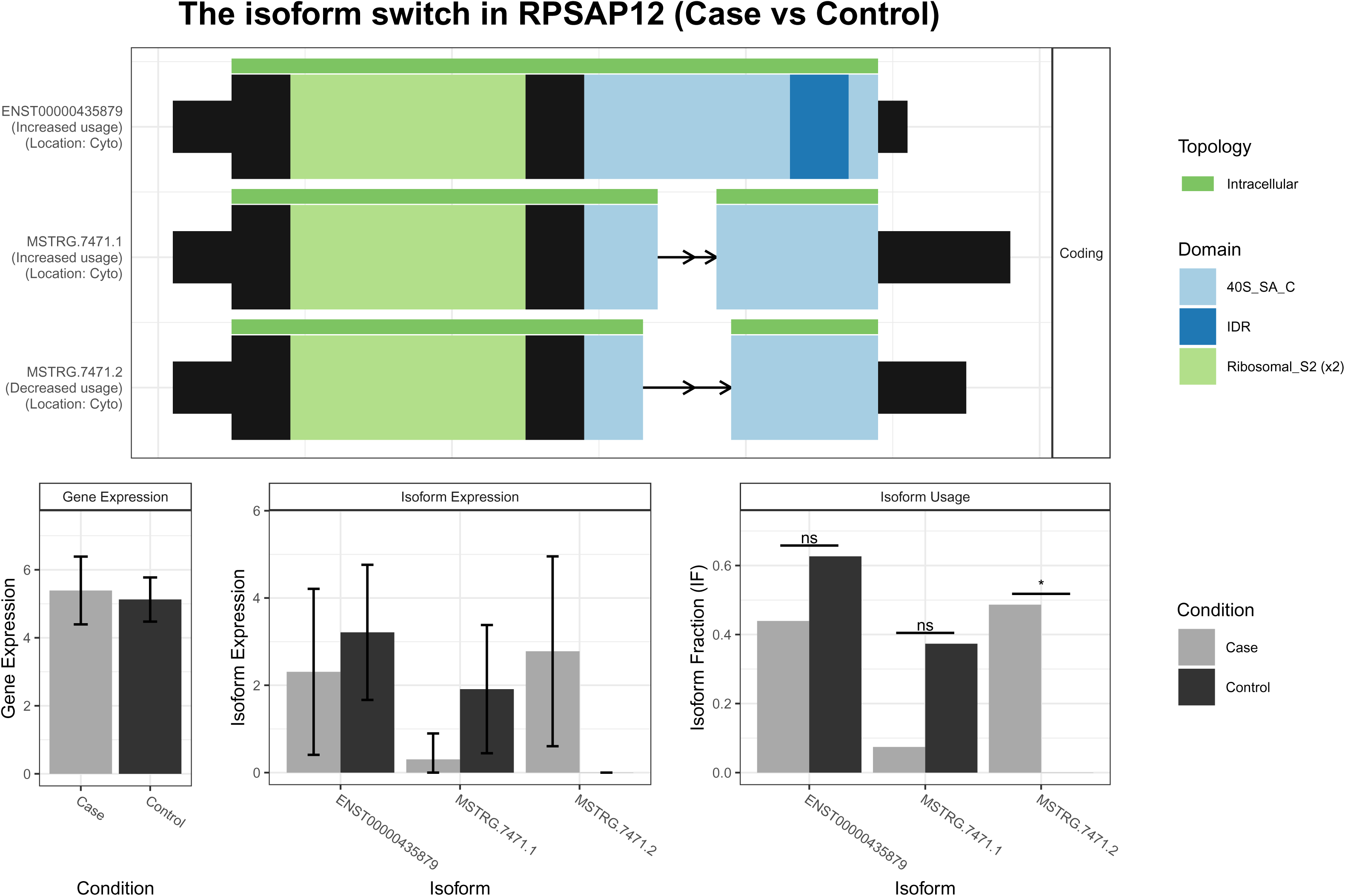

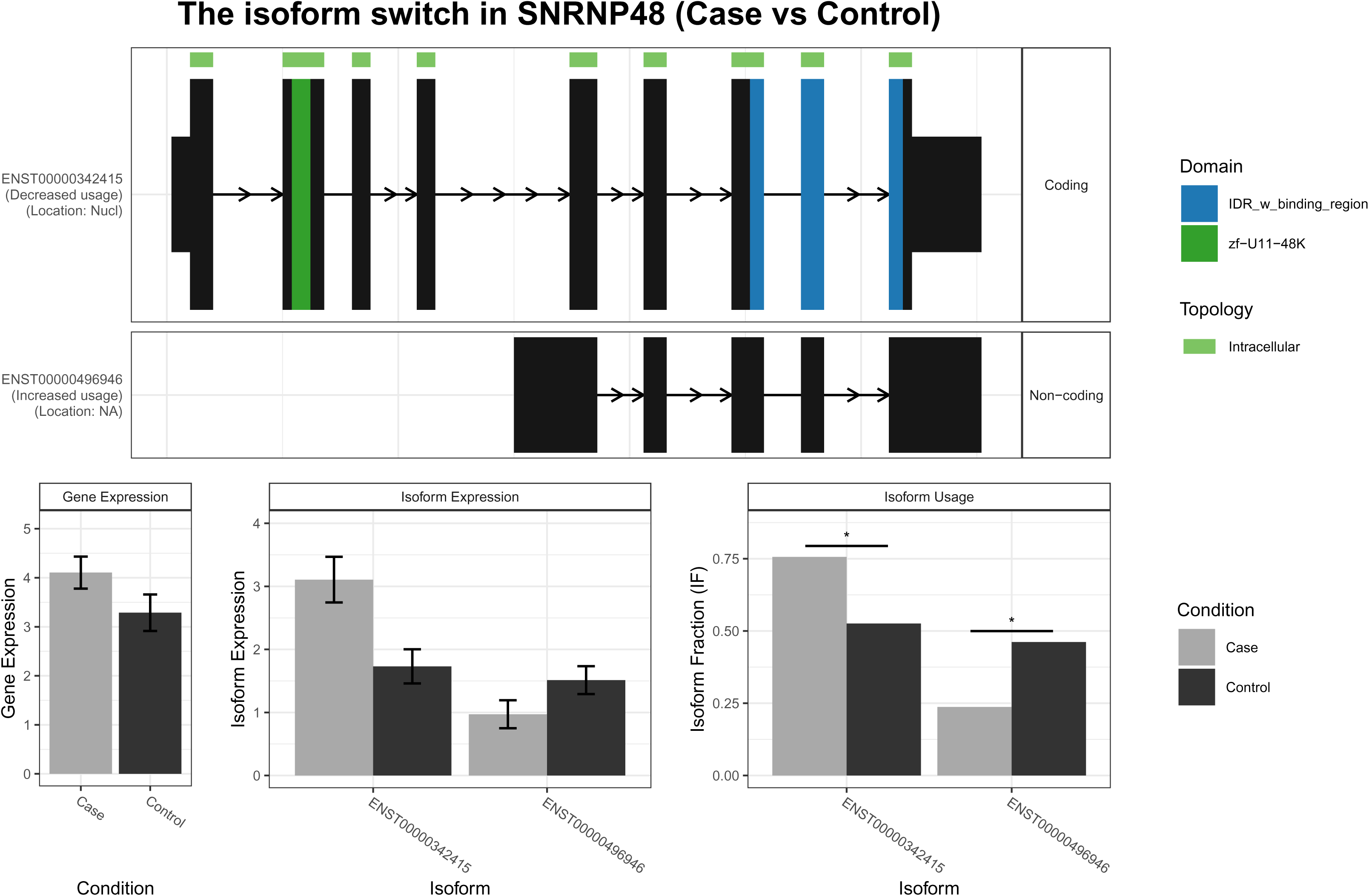

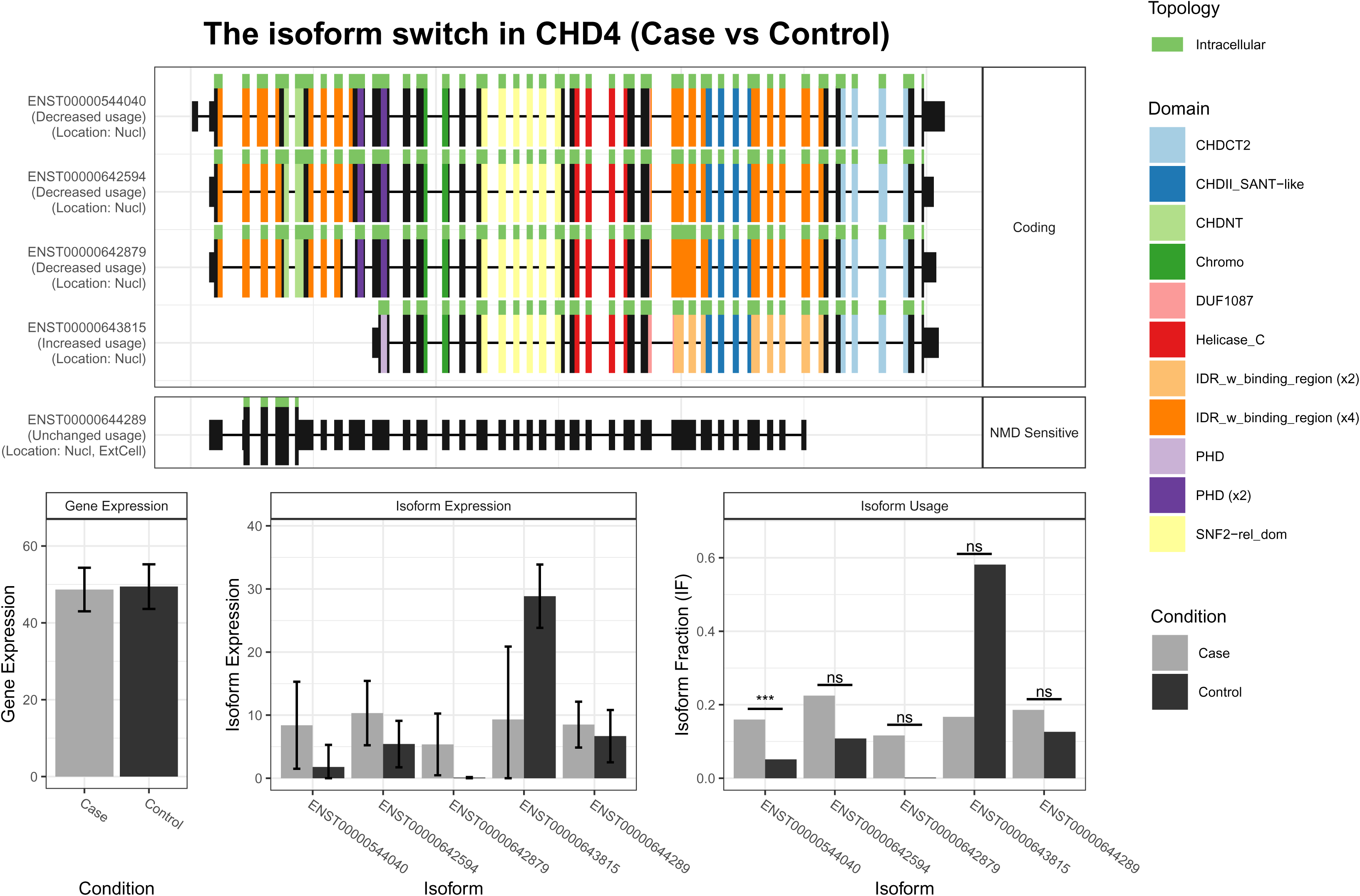

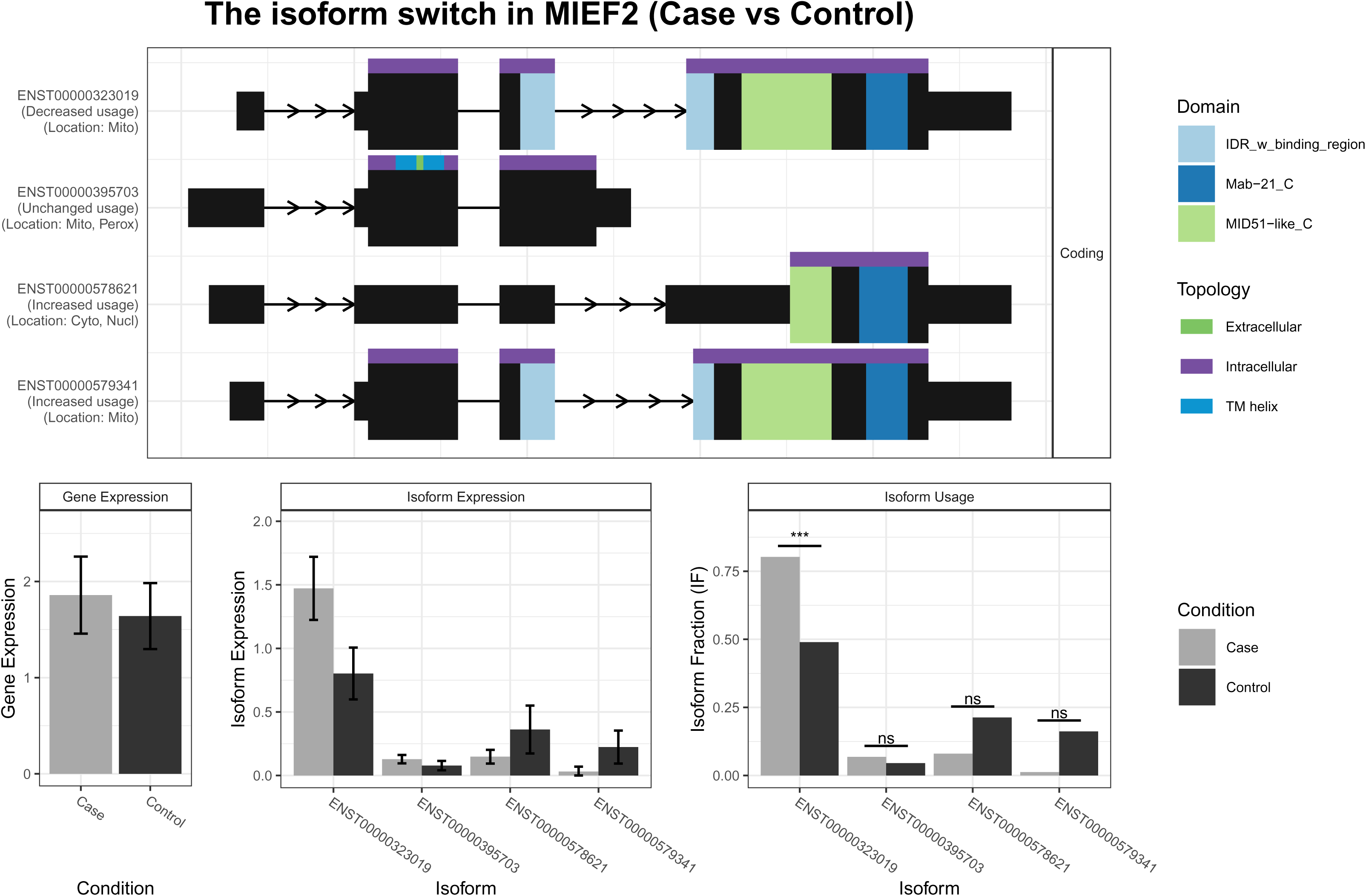

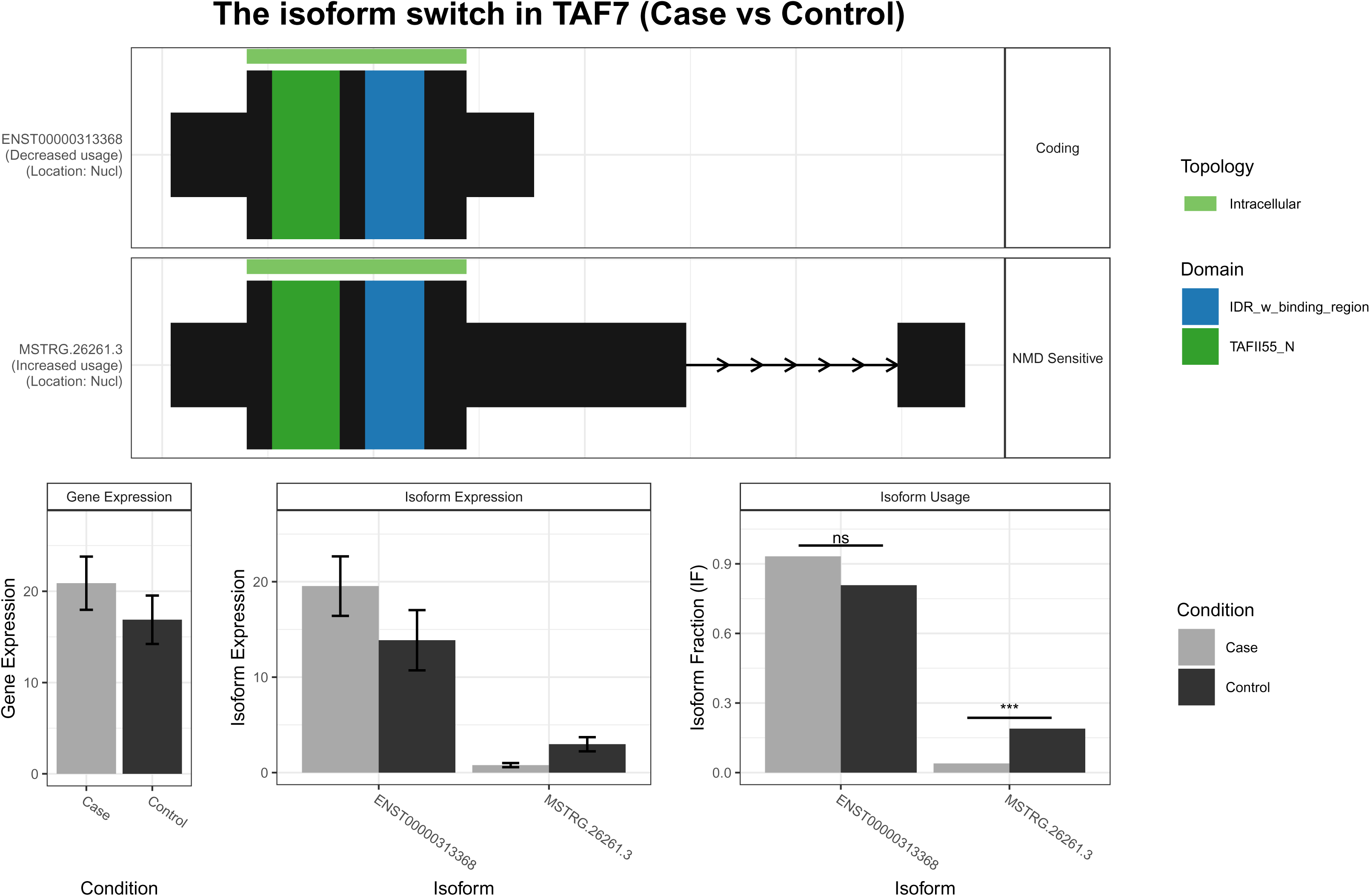

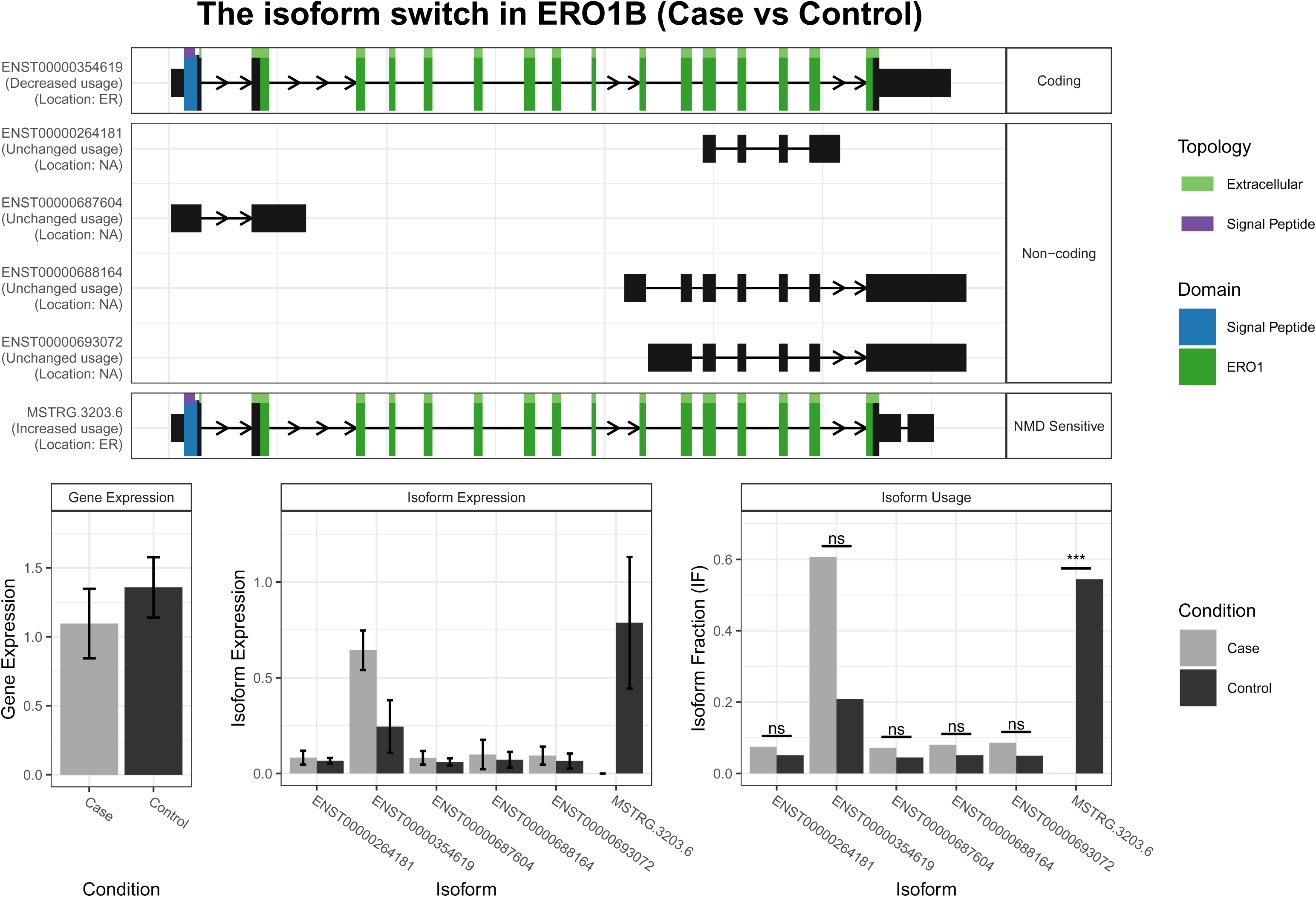

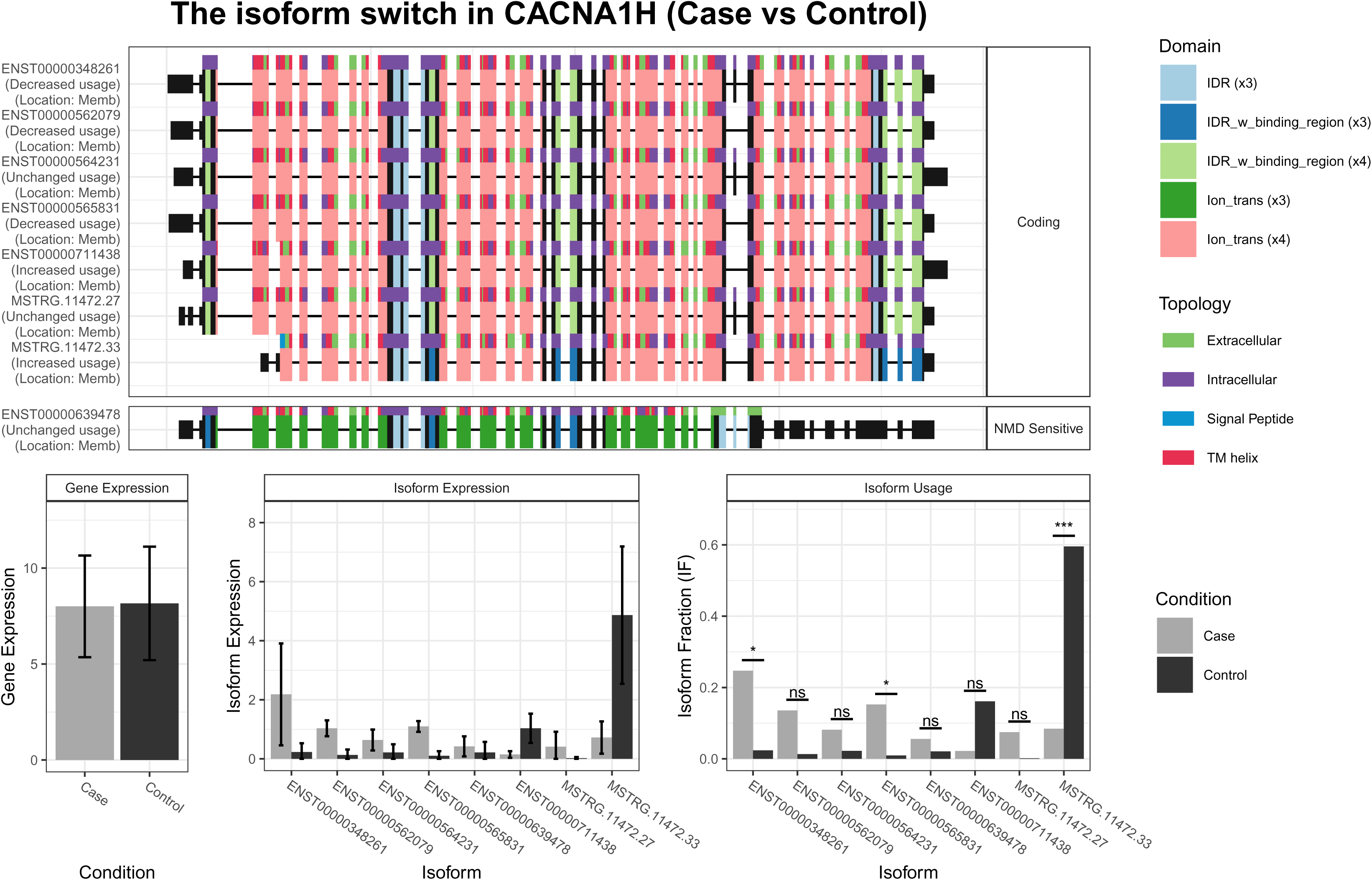

## References

1. Carruthers, BM, Anil Kumar Jain, Kenny L. De Meirleir, Daniel L. Peterson, Nancy G. Klimas, A. Martin Lerner, Alison C. Bested, et al (2003) Myalgic Encephalomyelitis/Chronic Fatigue Syndrome: Clinical Working Case Definition, Diagnostic and Treatment Protocols. Journal Of Chronic Fatigue Syndrome 11(1): p. 7–115. 10.1300/J092v11n01_02.

2. Sun, Y, Z Zhang, Q Qiao et al. (2024) Immunometabolic changes and potential biomarkers in CFS peripheral immune cells revealed by single-cell RNA sequencing. J Transl Med 22(1): p. 925. 10.1186/s12967-024-05710-w.

3. Todhunter-Brown, A, P Campbell, C Broderick et al. (2025) Recent research in myalgic encephalomyelitis/chronic fatigue syndrome: an evidence map. xx: p. xx. DOI: 10.3310/BTBD8846.

4. Wirth, KJ and C Scheibenbogen (2021) Pathophysiology of skeletal muscle disturbances in Myalgic Encephalomyelitis/Chronic Fatigue Syndrome (ME/CFS). J Transl Med 19(1): p. 162. 10.1186/s12967-021-02833-2.

5. Scheibenbogen, C and KJ Wirth (2025) Key Pathophysiological Role of Skeletal Muscle Disturbance in Post COVID and Myalgic Encephalomyelitis/Chronic Fatigue Syndrome (ME/CFS): Accumulated Evidence. J Cachexia Sarcopenia Muscle 16(1): p. e13669. 10.1002/jcsm.13669.

6. Walitt, B, K Singh, SR LaMunion et al. (2024) Deep phenotyping of post-infectious myalgic encephalomyelitis/chronic fatigue syndrome. Nat Commun 15(1): p. 907. 10.1038/s41467-024-45107-3.

7. Bizjak, DA, B Ohmayer, JL Buhl et al. (2024) Functional and Morphological Differences of Muscle Mitochondria in Chronic Fatigue Syndrome and Post-COVID Syndrome. International Journal of Molecular Sciences 25(3): p. 1675. 10.3390/ijms25031675

8. Wang, PY, J Ma, YC Kim et al. (2023) WASF3 disrupts mitochondrial respiration and may mediate exercise intolerance in myalgic encephalomyelitis/chronic fatigue syndrome. Proc Natl Acad Sci U S A 120(34): p. e2302738120. 10.1073/pnas.2302738120.

9. McCully, KK, BH Natelson, S Iotti et al. (1996) Reduced oxidative muscle metabolism in chronic fatigue syndrome. Muscle Nerve 19(5): p. 621–5. 10.1002/(SICI)1097-4598(199605)19:5<621::AID-MUS10>3.0.CO;2-Q

10. Steiner, S, A Fehrer, F Hoheisel et al. (2023) Understanding, diagnosing, and treating Myalgic encephalomyelitis/chronic fatigue syndrome - State of the art: Report of the 2nd international meeting at the Charite Fatigue Center. Autoimmun Rev 22(11): p. 103452. 10.1016/j.autrev.2023.103452.

11. Huang, K, BA Lidbury, N Thomas et al. (2025) Machine learning and multi-omics in precision medicine for ME/CFS. Journal of Translational Medicine 23(1): p. 68. 10.1186/s12967-024-05915-z.

12. Berk, M, JH Kim, LJ Williams et al. (2025) A novel discovery platform for targeted drug repurposing: application for psychiatric disorders. Lancet Psychiatry. 10.1016/S2215-0366(25)00066-5.

13. Bye, CR, E Qian, K Lim et al. (2026) Large-scale drug screening in iPSC-derived motor neurons from sporadic ALS patients identifies a potential combinatorial therapy. Nat Neurosci 29(1): p. 40–52. 10.1038/s41593-025-02118-7.

14. Williams, G, A Gatt, E Clarke et al. (2019) Drug repurposing for Alzheimer’s disease based on transcriptional profiling of human iPSC-derived cortical neurons. Transl Psychiatry 9(1): p. 220. 10.1038/s41398-019-0555-x.

15. Hargus, JJMPCG (2024) Induced pluripotent stem cell models as a tool to investigate andtest fluid biomarkers in Alzheimer’s disease andfrontotemporal dementia. Brain Pathology 34(4). 10.1111/bpa.13231.

16. Faried, A, R Rudiman, A Adam et al. (2026) Induced Pluripotent Stem Cells (iPSCs) as a Regenerative Strategy for Parkinson’s Disease: Opportunities and Challenges. Cell Biochemistry and Biophysics 84(2): p. 1679–1694. 10.1007/s12013-026-01993-6.

17. Cerneckis, J, H Cai, and Y Shi (2024) Induced pluripotent stem cells (iPSCs): molecular mechanisms of induction and applications. Signal Transduction and Targeted Therapy 9(1): p. 112. 10.1038/s41392-024-01809-0.

18. Duan, Y, DS Evans, RA Miller et al. (2020) signatureSearch: environment for gene expression signature searching and functional interpretation. Nucleic Acids Res 48(21): p. e124. 10.1093/nar/gkaa878.

19. Tanoli, Z, A Fernández-Torras, UO Özcan et al. (2025) Computational drug repurposing: approaches, evaluation of in silico resources and case studies. Nature Reviews Drug Discovery 24(7): p. 521–542. 10.1038/s41573-025-01164-x.

20. Dao Nguyen, TTTT, Bruna Panizzutti et al. (2026) iPSC-Derived Myogenic Progenitor Cells from People with ME/CFS Reveal Altered Genes and Pathways with Drug Repurposing Potential, 21 July 2026, . Research Square (PREPRINT). 10.21203/rs.3.rs-10015511/v2.

21. Chen, S, Y Zhou, Y Chen et al. (2018) fastp: an ultra-fast all-in-one FASTQ preprocessor. Bioinformatics 34(17): p. i884–i890. 10.1093/bioinformatics/bty560.

22. Kim, D, JM Paggi, C Park et al. (2019) Graph-based genome alignment and genotyping with HISAT2 and HISAT-genotype. Nat Biotechnol 37(8): p. 907–915. 10.1038/s41587-019-0201-4.

23. Liao, Y, GK Smyth, and W Shi (2014) featureCounts: an efficient general purpose program for assigning sequence reads to genomic features. Bioinformatics 30(7): p. 923–30. 10.1093/bioinformatics/btt656.

24. Yin, H, H Duo, S Li et al. (2025) Unlocking biological insights from differentially expressed genes: Concepts, methods, and future perspectives. Journal of Advanced Research 76: p. 135–157. 10.1016/j.jare.2024.12.004.

25. Leek, JT, WE Johnson, HS Parker et al. (2012) The sva package for removing batch effects and other unwanted variation in high-throughput experiments. Bioinformatics 28(6): p. 882–3. 10.1093/bioinformatics/bts034.

26. Yu, G, LG Wang, Y Han et al. (2012) clusterProfiler: an R package for comparing biological themes among gene clusters. Omics 16(5): p. 284–7. 10.1089/omi.2011.0118.

27. Shannon, P, A Markiel, O Ozier et al. (2003) Cytoscape: a software environment for integrated models of biomolecular interaction networks. Genome Res 13(11): p. 2498–504. 10.1101/gr.1239303.

28. Pertea, M, GM Pertea, CM Antonescu et al. (2015) StringTie enables improved reconstruction of a transcriptome from RNA-seq reads. Nature Biotechnology 33(3): p. 290–295. 10.1038/nbt.3122.

29. Wang, L, HJ Park, S Dasari et al. (2013) CPAT: Coding-Potential Assessment Tool using an alignment-free logistic regression model. Nucleic Acids Research 41(6): p. e74–e74. 10.1093/nar/gkt006.

30. Mistry, J, S Chuguransky, L Williams et al. (2021) Pfam: The protein families database in 2021. Nucleic Acids Research 49(D1): p. D412–D419. 10.1093/nar/gkaa913.

31. Teufel, F, JJ Almagro Armenteros, AR Johansen et al. (2022) SignalP 6.0 predicts all five types of signal peptides using protein language models. Nature Biotechnology 40(7): p. 1023–1025. 10.1038/s41587-021-01156-3.

32. Thumuluri, V, JJ Almagro Armenteros, Alexander R Johansen et al. (2022) DeepLoc 2.0: multi-label subcellular localization prediction using protein language models. Nucleic Acids Research 50(W1): p. W228–W234. 10.1093/nar/gkac278.

33. Hallgren, J, KD Tsirigos, MD Pedersen et al. (2022) DeepTMHMM predicts alpha and beta transmembrane proteins using deep neural networks. bioRxiv: p. 2022.04.08.487609. 10.1101/2022.04.08.487609.

34. Erdős, G, M Pajkos, and Z Dosztányi (2021) IUPred3: prediction of protein disorder enhanced with unambiguous experimental annotation and visualization of evolutionary conservation. Nucleic Acids Research 49(W1): p. W297–W303. 10.1093/nar/gkab408.

35. Vitting-Seerup, K and A Sandelin (2019) IsoformSwitchAnalyzeR: analysis of changes in genome-wide patterns of alternative splicing and its functional consequences. Bioinformatics 35(21): p. 4469–4471. 10.1093/bioinformatics/btz247.

36. Gilis, J, K Vitting-Seerup, K Van den Berge, et al. (2021) satuRn: Scalable analysis of differential transcript usage for bulk and single-cell RNA-sequencing applications. F1000Res 10: p. 374. 10.12688/f1000research.51749.2.

37. Van Booven, DJ, J Gamer, A Joseph, et al. (2023) Stress-Induced Transcriptomic Changes in Females with Myalgic Encephalomyelitis/Chronic Fatigue Syndrome Reveal Disrupted Immune Signatures. Int J Mol Sci 24(3).DOI: 10.3390/ijms24032698.

38. Vu, LT, F Ahmed, H Zhu et al. (2024) Single-cell transcriptomics of the immune system in ME/CFS at baseline and following symptom provocation. Cell Reports Medicine 5(1). 10.1016/j.xcrm.2023.101373.

39. Amir Shaghaghi, M, B Murphy, and P Eck (2016) The SLC2A14 gene: genomic locus, tissue expression, splice variants, and subcellular localization of the protein. Biochem Cell Biol 94(4): p. 331–5. 10.1139/bcb-2015-0089.

40. Bei, L, C Shah, H Wang et al. (2012) β-Catenin Activates the HOXA10 and CDX4 Genes in Myeloid Progenitor Cells. Journal of Biological Chemistry 287(47): p. 39589–39601.

41. Peng, Y, J Zhu, Q Zhang et al. (2024) Endogenous retroviral ERVH48-1 promotes human urine cell reprogramming. Cell Regen 13(1): p. 17. 10.1186/s13619-024-00200-2.

42. Nakka, K, C Ghigna, D Gabellini et al. (2018) Diversification of the muscle proteome through alternative splicing. Skelet Muscle 8(1): p. 8. 10.1186/s13395-018-0152-3.

43. Hinkle, ER, HJ Wiedner, AJ Black et al. (2019) RNA processing in skeletal muscle biology and disease. Transcription 10(1): p. 1–20. 10.1080/21541264.2018.1558677.

44. Bijlenga, P, JH Liu, E Espinos et al. (2000) T-type alpha 1H Ca2+ channels are involved in Ca2+ signaling during terminal differentiation (fusion) of human myoblasts. Proc Natl Acad Sci U S A 97(13): p. 7627–32. 10.1073/pnas.97.13.7627.

45. Li, S, M Hao, B Li et al. (2020) CACNA1H downregulation induces skeletal muscle atrophy involving endoplasmic reticulum stress activation and autophagy flux blockade. Cell Death & Disease 11(4): p. 279. 10.1038/s41419-020-2484-2.

46. Gehlert, S, W Bloch, and F Suhr (2015) Ca2+-Dependent Regulations and Signaling in Skeletal Muscle: From Electro-Mechanical Coupling to Adaptation. International Journal of Molecular Sciences 16(1): p. 1066–1095.

47. Bartsakoulia, M, A Pyle, D Troncoso-Chandía et al. (2018) A novel mechanism causing imbalance of mitochondrial fusion and fission in human myopathies. Hum Mol Genet 27(7): p. 1186–1195. 10.1093/hmg/ddy033.

48. Yamasaki, Y, K Sakamoto, S Yashiro et al. (2026) SMAD2 ubiquitination through PY motif regulates skeletal muscle mass and fibrotic degeneration. Scientific Reports 16(1): p. 6666. 10.1038/s41598-026-37582-z.

49. Gómez-del Arco, P, E Perdiguero, Paula S Yunes-Leites et al. (2016) The Chromatin Remodeling Complex Chd4/NuRD Controls Striated Muscle Identity and Metabolic Homeostasis. Cell Metabolism 23(5): p. 881–892. 10.1016/j.cmet.2016.04.008.

50. Klockner, I, C Schutt, T Gerhardt et al. (2022) Control of CRK-RAC1 activity by the miR-1/206/133 miRNA family is essential for neuromuscular junction function. Nature Communications 13(1): p. 3180. 10.1038/s41467-022-30778-7.

51. Giloteaux, L, KA Glass, A Germain et al. (2024) Dysregulation of extracellular vesicle protein cargo in female myalgic encephalomyelitis/chronic fatigue syndrome cases and sedentary controls in response to maximal exercise. J Extracell Vesicles 13(1): p. e12403. 10.1002/jev2.12403.

52. Petter, E, C Scheibenbogen, P Linz et al. (2022) Muscle sodium content in patients with Myalgic Encephalomyelitis/Chronic Fatigue Syndrome. J Transl Med 20(1): p. 580. 10.1186/s12967-022-03616-z.

53. Kuznyetsova, A and DA Hood (2025) Mitophagy in skeletal muscle: Impact of ageing, exercise and disuse. Exp Physiol. 10.1113/ep093041.

54. Tomas C, EJ, Strassheim V, Newton JL, Walker M (2020) The effect of myalgic encephalomyelitis/chronic fatigue syndrome (ME/CFS) severity on cellular bioenergetic function. PLoS ONE15(4): p. e0231136. 10.1371/journal.pone.0231136.

55. Appelman, B, BT Charlton, RP Goulding et al. (2024) Muscle abnormalities worsen after post-exertional malaise in long COVID. Nature Communications 15(1): p. 17.DOI: 10.1038/s41467-023-44432-3.

56. Agergaard, J, B Yamin Ali Khan, T Engell-Sørensen, et al. (2023) Myopathy as a cause of Long COVID fatigue: Evidence from quantitative and single fiber EMG and muscle histopathology. Clinical Neurophysiology 148: p. 65–75. 10.1016/j.clinph.2023.01.010.

57. Hejbøl, EK, T Harbo, J Agergaard et al. (2022) Myopathy as a cause of fatigue in long-term post-COVID- 19 symptoms: Evidence of skeletal muscle histopathology. Eur J Neurol 29(9): p. 2832–2841. 10.1111/ene.15435.

58. Mosegaard, S, G Dipace, P Bross et al. (2020) Riboflavin Deficiency-Implications for General Human Health and Inborn Errors of Metabolism. Int J Mol Sci 21(11). 10.3390/ijms21113847.

59. Schoenen, J, J Jacquy, and M Lenaerts (1998) Effectiveness of high-dose riboflavin in migraine prophylaxis. A randomized controlled trial. Neurology 50(2): p. 466–70. 10.1212/wnl.50.2.466.

60. Covarrubias, AJ, R Perrone, A Grozio et al. (2021) NAD(+) metabolism and its roles in cellular processes during ageing. Nat Rev Mol Cell Biol 22(2): p. 119–141. 10.1038/s41580-020-00313-x.

61. Dollerup, OL, S Chubanava, M Agerholm et al. (2020) Nicotinamide riboside does not alter mitochondrial respiration, content or morphology in skeletal muscle from obese and insulin-resistant men. J Physiol 598(4): p. 731–754. 10.1113/jp278752.

62. Nourbakhsh, B, N Revirajan, B Morris et al. (2021) Safety and efficacy of amantadine, modafinil, and methylphenidate for fatigue in multiple sclerosis: a randomised, placebo-controlled, crossover, double-blind trial. The Lancet Neurology 20(1): p. 38–48. 10.1016/S1474-4422(20)30354-9.

63. Minnich, A, N Tian, L Byan et al. (2001) A potent PPARα agonist stimulates mitochondrial fatty acid β-oxidation in liver and skeletal muscle. American Journal of Physiology-Endocrinology and Metabolism 280(2): p. E270–E279. 10.1152/ajpendo.2001.280.2.E270.

64. Coletta, DK, A Sriwijitkamol, E Wajcberg et al. (2009) Pioglitazone stimulates AMP-activated protein kinase signalling and increases the expression of genes involved in adiponectin signalling, mitochondrial function and fat oxidation in human skeletal muscle in vivo: a randomised trial. Diabetologia 52(4): p. 723–32. 10.1007/s00125-008-1256-9.

65. Wang, X, J Wang, C Ying et al. (2024) Fenofibrate alleviates NAFLD by enhancing the PPARα/PGC-1α signaling pathway coupling mitochondrial function. BMC Pharmacology and Toxicology 25(1): p. 7. 10.1186/s40360-023-00730-6.

66. Augustyniak, J, J Lenart, P Gaj et al. (2019) Bezafibrate Upregulates Mitochondrial Biogenesis and Influence Neural Differentiation of Human-Induced Pluripotent Stem Cells. Mol Neurobiol 56(6): p. 4346–4363. 10.1007/s12035-018-1368-2.

67. Ørngreen, MC, KL Madsen, N Preisler et al. (2014) Bezafibrate in skeletal muscle fatty acid oxidation disorders: a randomized clinical trial. Neurology 82(7): p. 607–13. 10.1212/wnl.0000000000000118.

68. Steele, H, A Gomez-Duran, A Pyle, et al. (2020) Metabolic effects of bezafibrate in mitochondrial disease. EMBO Molecular Medicine 12(3): p. EMMM201911589. 10.15252/emmm.201911589.

69. Bolaños, P and JC Calderón (2022) Excitation-contraction coupling in mammalian skeletal muscle: Blending old and last-decade research. Frontiers in Physiology 13 (2022). 10.3389/fphys.2022.989796.

70. David Olivares, VKD, Ying Shi, Debomoy K. Lahiri, Nigel H. Greig, Jack T. Rogers and Xudong Huang (2012) N-Methyl D-Aspartate (NMDA) Receptor Antagonists and Memantine Treatment for Alzheimer’s Disease, Vascular Dementia and Parkinson’s Disease. Current Alzheimer Research 9(6): p. 746–758. 10.2174/156720512801322564.

71. Liu, J, L Chang, Y Song et al. (2019) The Role of NMDA Receptors in Alzheimer’s Disease. Frontiers in Neuroscience 13 (2019). 10.3389/fnins.2019.00043.

72. Stoof, JC, J Booij, and B Drukarch (1992) Amantadine as N-methyl-D-aspartic acid receptor antagonist: new possibilities for therapeutic applications? Clin Neurol Neurosurg 94: p. S4–6. 10.1016/0303-8467(92)90006-o.

73. *NAMENDA (memantine hydrochloride) prescribing information*, FDA, Editor., 2018.

74. Generali, JA and DJ Cada (2014) Amantadine: multiple sclerosis-related fatigue. Hosp Pharm 49(8): p. 710–2. 10.1310/hpj4908-710.

75. Harandi, AA, H Pakdaman, A Medghalchi et al. (2024) A randomized open-label clinical trial on the effect of Amantadine on post Covid 19 fatigue. Scientific Reports 14(1): p. 1343. 10.1038/s41598-024-51904-z.

76. U.S. Food and Drug Administration (FDA). CINVANTI® (aprepitant) injectable emulsion, for intravenous use: Highlights of Prescribing Information. Revised March 2022. Available at: https://www.accessdata.fda.gov/drugsatfda_docs/label/2022/209296Orig1s008lbl.pdf. Accessed 1 October 2026.

77. Douglas, SD and SE Leeman (2011) Neurokinin-1 receptor: functional significance in the immune system in reference to selected infections and inflammation. Ann N Y Acad Sci 1217: p. 83–95. 10.1111/j.1749-6632.2010.05826.x.

78. Schank, JR and M Heilig (2017) Substance P and the Neurokinin-1 Receptor: The New CRF. Int Rev Neurobiol 136: p. 151–175. 10.1016/bs.irn.2017.06.008.

79. Yang, Y, W Zhou, X Xu et al. (2022) Aprepitant Inhibits JNK and p38/MAPK to Attenuate Inflammation and Suppresses Inflammatory Pain. Frontiers in Pharmacology 12 (2021). 10.3389/fphar.2021.811584.

80. Zhao, XN, ZZ Bai, CH Li et al. (2020) The NK-1R Antagonist Aprepitant Prevents LPS-Induced Oxidative Stress and Inflammation in RAW264.7 Macrophages. Drug Des Devel Ther 14: p. 1943– 1952. 10.2147/dddt.S244099.

81. Yasunari, K, K Maeda, M Nakamura et al. (2005) Benidipine, a long-acting calcium channel blocker, inhibits oxidative stress in polymorphonuclear cells in patients with essential hypertension. Hypertens Res 28(2): p. 107–12. 10.1291/hypres.28.107.

